# Grapevine vigour: a critical factor driving Trunk Disease expression

**DOI:** 10.1101/2024.06.11.598450

**Authors:** Marion Claverie, Maïko Audras, François Berud, Didier Richy

**Affiliations:** Institut Français de la Vigne et du Vin, Pôle Rhône-Méditerranée, Institut Rhodanien, 2260 Route du Grès, 84100 Orange, France; Chambre d’Agriculture du Gard, 1120 route St-Gilles, CS 38283, 30942 Nîmes Cedex 9, France; Chambre d’Agriculture de Vaucluse, 2260 Route du Grès, 84100 Orange, France; Chambre d’Agriculture des Bouches du Rhône, Maison des Agriculteurs, 22 Avenue Henri Pontier, 13626 Aix en Provence Cedex 01, France

**Keywords:** Grapevine, trunk diseases, esca, vigour, Grenache N, dieback, decline

## Abstract

Esca and Black Dead Arm (EBDA) are complex grapevine trunk diseases (GTD) that are major causes of mortality and production decline in French vineyards. Unravelling the different factors that determine symptom expression is crucial in mitigating the impact on growers. While cultivar, climate, age, and pruning system are known contributors, they do not fully explain the variability in EBDA occurrence. This study investigates the role of grapevine vigour as a determinant factor in EBDA expression.

In 2022 and 2023, three vineyard networks were monitored, each consisting of approximately 30 vineyard plots of Grenache Noir, uniform in age. To minimize climate variation the selected vineyards were located within small geographical areas. We evaluated grapevine vigour and its primary drivers—water and nitrogen status, weed cover, production, and vegetation biomass—and correlated these factors with EBDA occurrence.

Results show that current-year vigour is positively correlated with the EBDA incidence rate in all network*year scenarios. The relationship curve suggests that, while low to moderate vigour is consistently associated with reduced symptom expression, high vigour can be correlated with either high or low expression and implies the involvement of additional factors. In one instance, previous year water stress, of all tested variables, was most correlated with current year EBDA incidence, though vigour was also influential. In that case, EBDA expression seems to be maximal when water stress the year before is combined with a substantial current year spring vigour. While these results need to be confirmed over a longer period, in more regions and with other cultivars, they open new perspectives of applications for growers. They suggest a wider implication of grapevine physiology than just vigour.

## Introduction

Trunk diseases are a major cause of grapevine dieback worldwide (Guérin Dubrana et al., 2019; La Fuente et al., 2016). In France, it causes a heavy economic burden for wineries and vine growers, especially for the last 20 years since an increase in trunk disease expression has been observed in many winegrowing regions.

Among trunk diseases, esca and Black Dead Arm (BDA) cause foliar symptoms on leaves, shoots, and grapes up to complete drying and wilting of the vine and culminating in the death of the vine, generally within a few years. This disease is thought to be due to a complex of different fungi that enter the vine through wounds, mainly during winter pruning and shoot thinning in the springtime and develop inside the wood causing necrosis and decay. Starting from a necrosed vine, foliar symptoms may or may not appear and fluctuate from one year to another. Very few attempts in reproducing the typical symptoms from fungal inoculations have been successful, and for all these reasons, esca and BDA are considered to be multi-factorial complex diseases that still remain only partially understood (for reviews see Bertsch et al., 2013; Claverie et al., 2020; Larignon et al., 2009; Mugnai et al., 1999; Surico et al., 2006; Wagschal et al., 2008). As esca and BDA show a lot of similarities both in their symptom appearance and their supposed causal agents and mechanisms, they will be considered together in this article and referred to as EBDA.

In contrast with perennial inner symptoms, the external incidence of EBDA in vineyards is highly variable (Fernandez et al., 2024), depending on the year, the cultivar, the age of the vineyard (Etienne et al., 2024; Gastou et al., 2024; Grosman & Doublet, 2012), the pruning type and training management (Dal et al., 2008; Kraus et al., 2022; Lecomte et al., 2019; Travadon et al., 2016), and the climate (Bortolami et al., 2021; Calzarano et al., 2018; Larignon, 2009, 2020; Lecomte et al., 2023; Marchi et al., 2006; Surico et al., 2000). There is a big challenge in understanding what determines disease symptom expression, to try to mitigate the impact of the disease for growers. But even in the same region, in the same year, with the same cultivar and age, there are still big potential differences between vineyard plots (Dewasme et al., 2022; Etienne et al., 2024; Li et al., 2017; Monod et al., 2023). This incited us to explore why and to set our work at an even smaller scale than the previously mentioned studies, i.e. local networks of a few kms wide. At such a scale, climate is supposed to show little variation, and that was conducive to test the hypothesis of an influence of the vigour of the plot.

‘Vigour’ is a word that, at least for grapevine, encompasses many things: the vigour of a plant characterizes its ability to grow and produce vegetative biomass, from leaves to grapes, roots and wood. For grapevine, Champagnol (1984) states that vigour is the indicator of an intense metabolic activity of the growing plant that depends on external factors (mainly the environment) and grapevine factors (grapevine genetics, bud load and quality of sap flow, amount of reserve material etc). To quote from Reynier (2011), “*A vigorous grapevine has an active growth, shoots with long internodes, numerous lateral shoots, an elevated number of bunches*.” But Champagnol restricts the use of the word ‘vigour’ to the mean biomass of one shoot or cane, i.e. the total vegetative biomass per number of shoots. Vigour can sometimes also be applied to the potential of the environment, proper functioning and fertility of the soil and its ability to feed the vine and produce biomass. In this work, we considered ‘vigour’ as the expression of total vegetative biomass of a grapevine plant. Because grapevine vigour is closely related to many components of the environment, like climate (rain, temperature, VPD…), soil characteristics, genetic profile of the grapevine itself (cultivar, rootstock and clone), competition with other vines (planting density) or other plants (weeds in particular) and cultural practices (fertilization, irrigation, soil management…), we can be tempted to consider the system as a whole, including vine vigour and its determinants.

In the past decades of research on grapevine trunk diseases, there is not much scientific data about the relationship between grapevine vigour and EBDA: vigour is not mentioned as a determinant factor in most of the reviews on EBDA (Bertsch et al., 2013; Gramaje et al., 2018; Mondello et al., 2018; Mugnai et al., 1999) even if the influence of environmental factors on EBDA, particularly heat and water stress (Bortolami et al., 2021; Calvo-Garrido et al., 2021; Fischer & Kassemeyer, 2012; Fischer & Peighami-Ashnaei, 2019; Songy et al., 2019; van Niekerk et al., 2011), and conversely of EBDA on grapevine physiology (as reviewed by Fontaine et al. (2016)), have been studied and documented much more. Nevertheless, in the field, some empirical evidence of the influence of vigour sometimes emerges: vine growers or field advisors occasionally mention a positive vigour effect on EBDA or mortality, generally on plots showing a gradient of vigour, for example due to the effect of a slope or the competition with a border of nearby trees (as also reported by Surico et al. (2000)). In that case, vines located in the higher competition zone are less vigorous and show less symptoms of EBDA. To illustrate this point, an assessment of EBDA incidence was done in 2023 in a commercial 19-year- old vineyard plot of cv. Mourvèdre located in the Rhone Valley, where a hedge of shrubs had been planted on one side parallel to the vine row one year before planting the vineyard. The first 4 rows, that proved to be most in competition with the hedge, were significantly less affected by EBDA symptoms than control rows (located 25m from the first row), with an average of 9% and 19% symptomatic vines, respectively (personal communication).

Apart from empirical observations, some scientific references on the influence of vigour exist: between 2004 and 2006, Destrac-Irvine et al. (2007) monitored a network composed of 22 vineyard plots located in different sub-regions of Bordeaux. They observed that, beside an overall decrease in EBDA expression on the driest year (2005), incidence of EBDA in a given year was higher in plots with a higher water holding capacity, comfortable nitrogen status and yield (Lecomte et al., 2011). In addition, in a long-term trial comparing different ground vegetation cover and N fertilization combinations in the Cognac region of France, Dumot (2022) assessed mortality rates after 20 years of study. Treatments consisted of vineyard blocks with different doses of fertilizers and levels of ground vegetation cover. Mortality, caused mainly by EBDA, was highest in the treatment with higher fertilization (60 N unit per ha) and no ground vegetation cover and was lowest in the non-fertilised block with ground vegetation cover. Kuntzmann et al. (2013) investigated the role of a set of cultural factors in explaining the expression of EBDA on a network of plots of different cultivars and ages monitored between 2003 and 2011 in the Alsace region of France. Among the different significant factors that were pointed out, these authors mentioned vigour and yield were frequently associated with higher expressions of EBDA, even if he also points out discrepancies with other studies and years where higher EBDA expression was associated with lower vigour. Lately, in some more recent work on the Alsacian network, higher incidence rates have been associated with plots showing lower soil water holding capacities (Abidon et al., 2019). Finally, two recent studies included vigour and their related variables among the factors tested for their influence on EBDA incidence: Monod et al. (2023) showed that, on a network of plots of the same cultivar (Gamaret), age and planting material origin set among 4 Swiss wine producing regions, the variables most correlated to EBDA incidence were climatic variables (especially rainfall in May and June) and soil water holding capacity; vine vigour being not significant, though showing a slight correlation with EBDA symptom expression and vine mortality. In the study conducted by Gastou et al. (2024), consisting of 46 cultivars in one vineyard plot in Bordeaux in which monitoring was done on epidemiological and biotic variables (vine vigour, nitrogen and δC13 indices, phenology), water use efficiency of the cultivar, and to a lesser extent vigour, proved to be both positively correlated to EBDA incidence and mortality.

In this study, we intended to test if vine vigour, in relation with its main drivers, could be related to EBDA symptom expression during a 2-year course of monitoring. To do so, we worked on 3 small local networks of vineyard plots of the same vine cultivar Grenache N with similar ages and no or little climatic variation between plots during the same year.

## Materials and methods

### 1. Three geographical networks each composed of ∼30 vineyard plots

This work was conducted in south-eastern mediterranean wine regions of France (regions *Sud* and *Occitanie*, France) in 3 different networks of vineyard plots located in the Bouches-du-Rhône (named *Network-B, 43.6537° N, 5.2624° E*), Gard (named *Network-G, 44.1028°N, 4.6578°E*) and Vaucluse (named *Network-V, 44.268 7° N, 5.1252 ° E*) departments. Each network is composed of 31, 26 and 25 plots for Network-B, *Network-G* and Network-V respectively, and covers a small geographical area of a few kilometres in diameter, corresponding to a group of 3 to 6 nearby towns.

The plots are productive commercial vineyards and belong to 58 different vine growers. All plots of a given network are part of the same “cooperative” winery.

All plots are planted with the same cultivar, Grenache Noir, between 16 and 26 years old in 2021 (median age of 20.5 years, 55% of the plots between 18 and 22 years old). Network-B is mostly aimed at producing rosé wines (yield objective of 60 hl per ha), while the other two produce red Côtes-du- Rhône wines (yield objective of 40 to 50 hl per ha). Vine planting densities are quite similar between plots, from 3500 to 4500 vines per hectare, with an average around 4000.

For each network, the selection of the vineyard plots monitored in 2022 and 2023 is a subsample deriving from a larger 100-plot network where a preliminary trunk disease incidence evaluation was conducted in 2021. The selection criteria were chosen in order to (1) keep all the plots showing maximum and minimum incidence of EBDA in 2021, as well as intermediate levels, (2) tighten vine ages around 20 years-old (which has shown to be the age of maximum EBDA incidence) to minimize a bias effect of age and (3) ensure the selected subsample properly reflects the diversity of agronomical situations in the area (in terms of irrigation and soil management in particular).

### 2. Agronomical characteristics of the three networks

All 3 networks have a mediterranean climate (average Huglin index 2400-2600°C over the last decade; average wetness soil index: -100 to -180mm between 2017 and 2023). The altitudes vary from 170m to 370m in Network-B, 30m to 200m in Network-G and 260m to 450m in Network-V.

The principal soil types found in the study area are clayey calcareous soils (*calcosols*), loamy and sandy soils from colluvial deposits (*colluviosols*) and, in a few cases, soils from alluvial deposits as well as particular soil types like chromic soils (*fersialsols)* and cambisols (*brunisols)*.

In Network-B, 16 plots are irrigated and 14 are not. Soil management is mostly tillage both under the row and in the inter-row with generally little weed cover remaining. Only 4 plots out of 30 are intentionally covered with spontaneous or sown species in the inter-row.

Network-G, on the other hand, is mostly non irrigated (only 3 plots are irrigated) and the soil is most frequently covered with a permanent natural cover in the inter-row, for half of the plots in all inter- rows, and the other half only in one inter-row out of two, sometimes less. Only 4 out of 26 plots are tilled in both rows and inter-rows.

Finally, Network-V is not irrigated, and only 7 out of 25 plots have natural ground cover in the inter- rows, the rest are tilled.

In plots where the soil management is tillage, it is important to note that some plots may have no ground cover whereas others may still show substantial amounts of weed cover during the year.

The main training system is spur-pruned double cordon, with only one plot goblet trained, and one with mechanical pruning.

The rootstock is known for only 75% of the plots; the most common one is 110R (63% of the known rootstocks), followed by 140Ru (27%).

The plots can be affected by other dieback diseases than EBDA, such as the fan leaf virus or root rot due to *Armillaria*. Eutypa dieback and ‘yellows’ phytoplasma diseases remain scarce, with only a few plots and a few vines per plot affected.

### 3. Monitored variables during 2021, 2022 and 2023

Variables monitored in this study are water uptake, nitrogen status and grapevine vigour, as well as ground vegetation cover. Grapevine yield and vegetation were also estimated. Climatic data was collected, as well as EBDA incidence. The different variables and indicators used are shown in table1.

The **climate data** (temperature, rain and VPD) used derive from Antilope and Arome models provided by *Météo France* using a one-km grid of the territory. These spatialized data are calculated integrating different data sources (real weather stations, satellite images and RADARs). Climate estimations can be done on any location using the 4 nearest points of the grid.

Two indicators describe the **water uptake and status of the vine:** (1) Visual observation of the slowing down of shoot growth using the “apex method” (Pichon et al., 2022). At each measurement date, 50 apical meristems are assessed depending on their growth profile (1=full growth; 0.5=moderate growth and 0=stopped growth). The IC growth index is then calculated (ICapex) using the following formula:

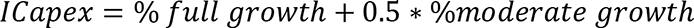

Where % full growth is the number of apexes in full growth out of the 50 apexes assessed. As we can see from the formula, the higher the value of ICapex, the less the water stress. Assessment is made at 2 or 3 different stages during the summer between berry growth and ripening, generally around late June (berry growth), late July (early veraison) and late August (ripening). To integrate the 3 dates into one indicator, the Area Under the Curve (AUC) of ICapex was calculated as follows:

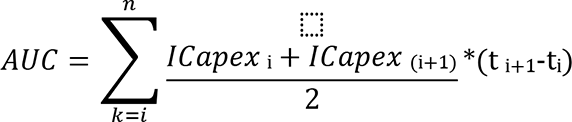

Where ti is the Julian day of the measure i. As for ICapex, the higher the value of AUC, the less the water stress. In 2023, an additional assessment was done at flowering time, to characterize spring vigour. This earlier measure was not added to the AUC calculation, so that the AUC values in 2022 and 2023 are done on the same basis and remain comparable. (2) Carbon isotope discrimination analysis (δC13) (Gaudillère et al., 2001) was done in 2022 on only half of the plots using a 200-berry sample that was randomly picked at harvest time. On the same day, a rapid visual assessment of the water status of the vine at harvest was done using grades: 0 if shoot growth is still active, 1 if growth has ceased and 2 if drought symptoms were present (basal leaves drying and fall).

The **chlorophyll index** is used to account for **nitrogen status of the vine**. The N-Tester (Yara, Oslo, Norway) or SPAD 502 (Konica Minolta, Nieuwegein, Netherlands) devices were used depending on the network. On each plot, chlorophyll index is given after 30 measurements (pinching of the leaves in the fruit zone) distributed up and down along 2 rows. The measurement was performed at veraison time, except for Network-G in 2022 (pre harvest time). In 2023, an additional measurement was performed in May at pre-flowering time, along with spring vigour characterization. At harvest time, in addition to the visual assessment of water status of the vine, a rapid visual assessment of nitrogen status of the canopy was done as well, based on aspect of the canopy (intensity of green colour and size and number of the leaves): 0 if nitrogen deficiency (typical pale light green canopy), 1 if moderate green colour and 2 if dark green and/or large basal leaves indicating high nitrogen nutrition.

**The ground vegetation cover** evaluation was done visually, using a 4-grade scale based on the percentage of the ground covered by weeds or crops (from 0=no ground vegetation to 4=entire surface covered by ground vegetation), and discriminating the row from the inter-rows (whether tilled or not), as well as discriminating the green living part (considered as the active weed cover part) from the total covered area. The ground cover was assessed at the same dates as the apex growth. Different indexes can then be used to characterize a plot depending on those different criteria and the date of the measurement. These indexes are generally more or less inter-correlated within the same year, and only some of them have been kept in this article, i.e. index characterizing the row cover, the average annual score on (row + inter-row) and the average proportion of green active cover during the season.

**Grape production** and **vegetative biomass** are visually assessed at harvest time, like water and nitrogen status, using visual scoring. The assessment of production is based on the achievement of the yield objective for the considered network: 1 if production is far under the objective, 2 if production is under the objective, 3 if production reaches the objective and 4 if it is beyond. For vegetation, it is a subjective expert note based on the volume and density of the canopy, from 1 (very small) to 4 (big vegetation). Two operators proceed and compare their score. These scoring methods have been used because of their good (time consuming/robustness) ratio observed in previous studies.

**Vigour assessment** was done in winter, by counting the number of canes (CN) on 20 vines per plot and measuring cane basal diameter (d) on about 30 to 40 canes per plot in order to calculate the section area (SA) of the canes. Cane diameter was measured using an electronic calliper (precision 0.1mm), on all canes of one spur per vine on half of the 20 vines, between inter-node 2 and 4 at the base of the cane. When possible, the approximate height (H) of the canopy hedge was measured as well. Therefore, vigour proxy that was used is CN*SA (CN*π*d²/4). In 2022, vigour assessment has only been done on part of the plots (23 out of 30, 16 out of 29 and 13 out of 25 respectively in Network-B, 30 and Network-V). On 13 out of 23 plots of Network-B, the plots had been previously pruned by the growers. In that case, diameter measurement was done on canes collected on the ground (entire canes i.e. including the first inter-node) and cane number by counting the number of spurs left on the vine and applying a correction factor of 2.6, which corresponds to the average number of canes per spur counted on the remaining unpruned plots. In this article, “CN*SA” is frequently called “vigour” proxy even if it is *sensu stricto* a proxy of total vegetative biomass the vine.

In order to better characterize spring vigour conditions, measurements of water and N status have been added to the protocol in 2023. At pre-flowering time, i.e. stage 9-12 detached leaves (BBCH 57) for Network-G, Network-V and mid-flowering (BBCH 60-65) for Network-B, we measured (1) growth status of the apex (‘apex method’) and (2) chlorophyll index, using the same protocol as for water and N status assessment (see above). As a probable bias of the phenological stage (between BBCH 57 and flowering) was visible on chlorophyll index (higher average values in Network-B measured at flowering than on the other 2 networks measured before flowering, *not shown*), we used the normalized values per network.

Irrigation schedules and amounts were collected from growers at the end of the season. In addition to characterizing the climate, climate data were also used to calculate water balance (WB) indexes. We followed (Carbonneau, 1994) simplified method for theoretical (WBth) and estimated water balance (WBest). In both cases, soil water holding capacity was fixed at 200mm and irrigation doses were included in addition to rainfall amounts. In WBest only, the ground cover is considered using % of full surface covered for its contribution to soil transpiration. In 2023, the precipitation regime was chaotic in the spring sometimes inducing variability inside a given network, rainfall data used to run WB were localized: all plots have thus been affected to a subregion of the network where the rainfall amounts were homogeneous. In 2022, only one median plot per network was used to collect rainfall data because of very homogeneous conditions in each network. Consequently, for a given network and year where very little variation in climate data is observed, in the absence of irrigation, WBest is mainly reflecting ground vegetation cover (R²>0.90, *not shown*).

Finally, the assessment of **EBDA incidence** was conducted prior to harvest (i.e. from the end of August to mid-September), on 200 to 600 living vines per plot (average 350-400 depending on the network), in at least 4 rows per plot, in order to represent the total surface of the plot using the same fixed row numbers over the three years. EBDA well trained operators counted the number of symptomatic vines, discriminating apoplexy form from the typical EBDA symptoms, as represented in supplementary figure 1. Typical EBDA form of symptoms is “tiger-stripe-like” mild form of symptoms on the leaves (supplementary figure 1 a). But cultivar Grenache N is particularly prone to show severe form of EBDA (supplementary figure 1 b&c), with frequent leaf drying and falling, sometimes affecting only a few leaves on a shoot, but more frequently entire shoots and vines. Besides, apoplexy form can also be regularly seen; it corresponds to a sudden wilting and drying of the entire vegetation (sometimes only half a vine, but always affecting more than a shoot) with a homogeneous aspect between leaves and shoots (supplementary figure 1d, resembling as if the vine trunk had been ploughed off), oppositely to severe EBDA form where shoots are affected to different extents and remaining green parts can still be seen.

### 3. Statistical analysis

All statistical analyses were conducted using XLStats software (Lumivero, Paris, France): correlation matrix using Pearson and Spearman tests, PCA and Kruskal-Wallis non parametrical tests (threshold p=0.05 except when specified).

## Results

### 1. Characteristics of the vineyard networks: EBDA, climate and agronomical indicators

#### 1.1. Differential expression of EBDA symptoms per network over the 3 year-period

The average values of typical EBDA and apoplexy for each network and year are shown on figure 1a. Total EBDA ranges between 1.8% and 6.1%. Notwithstanding the important deviation observed inside each network*year, which was expected, EBDA expression show some significant differences between networks and years (Network-G in 2021 with greatest incidence and Network-G and Network-V in 2022 showing least).

**Figure 1.**
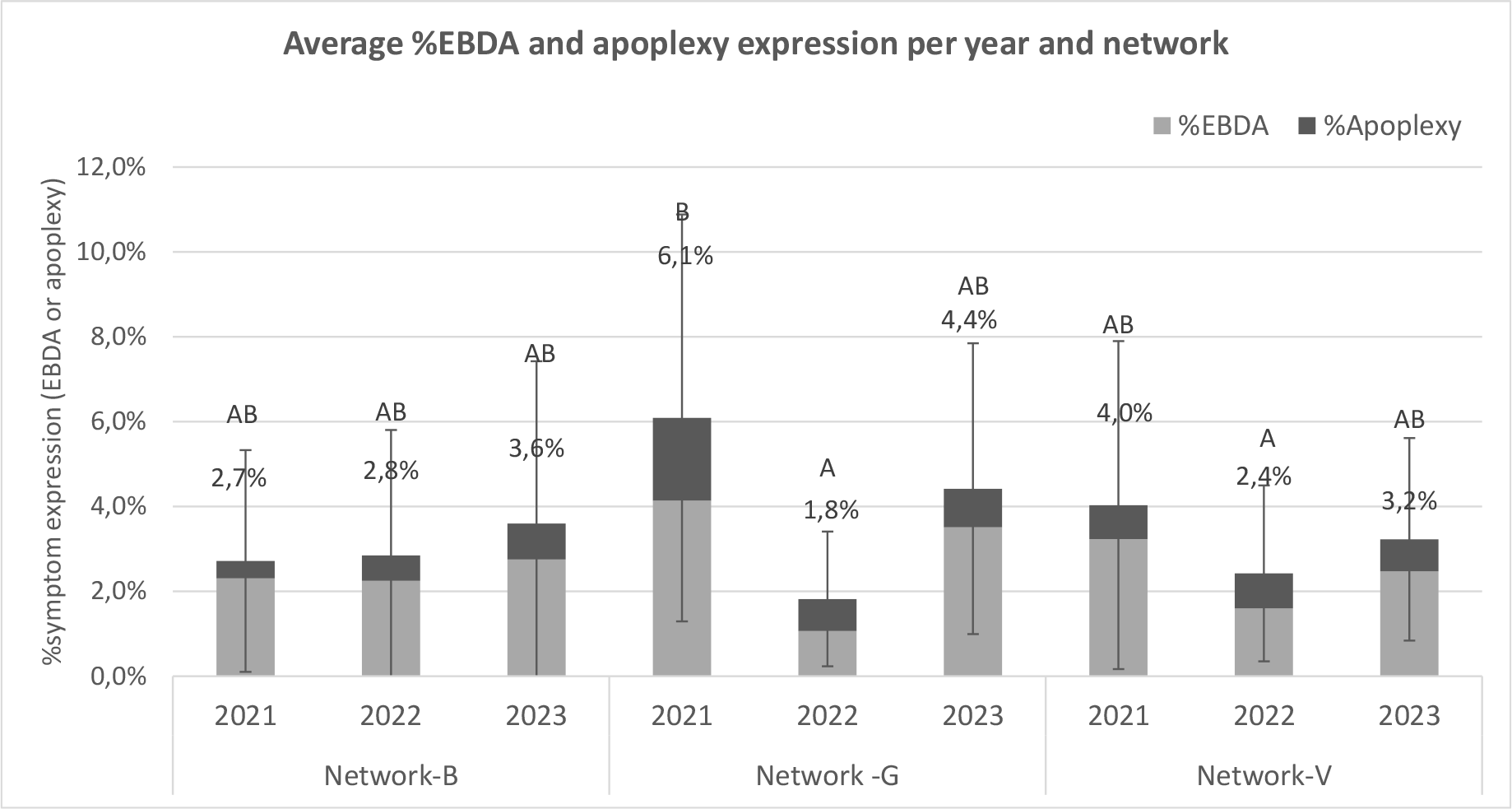
Mean values of EBDA and apoplexy per year for each local plots Network-B, Network-V and Network-G. Bars=standard deviation; letter= Kruskal-Wallis nonparametric test groups

Figure 2 illustrates the diversity of EBDA incidences among networks and years. The network- dependant year effect highlighted on figure 1 is manifest on these graphs: for Network-B, almost no year effect is visible on the EBDA total expression between 2021, 2022 and 2023 (all plots in line close to the bisector). For Network-V, 2022 expression is reduced by a factor of 2 when 2023 expression reaches 80% of its 2021 level, both years showing similar ranking between plots as 2021. For Network-G in contrast, the expression rate in 2022 is reduced by a factor of 4 compared to 2021, but the ranking of the plots is quite conserved, contrarily to 2023 expression, which is higher than 2022 and slightly lower than 2021, but with a ranking of the plots that seems to be mixed up. This change in ranking positions between plots between 2021 and 2023 is not due to apoplexy, as the graph looks the same when made with typical EBDA symptoms alone (*not shown*).

**Figure 2.**
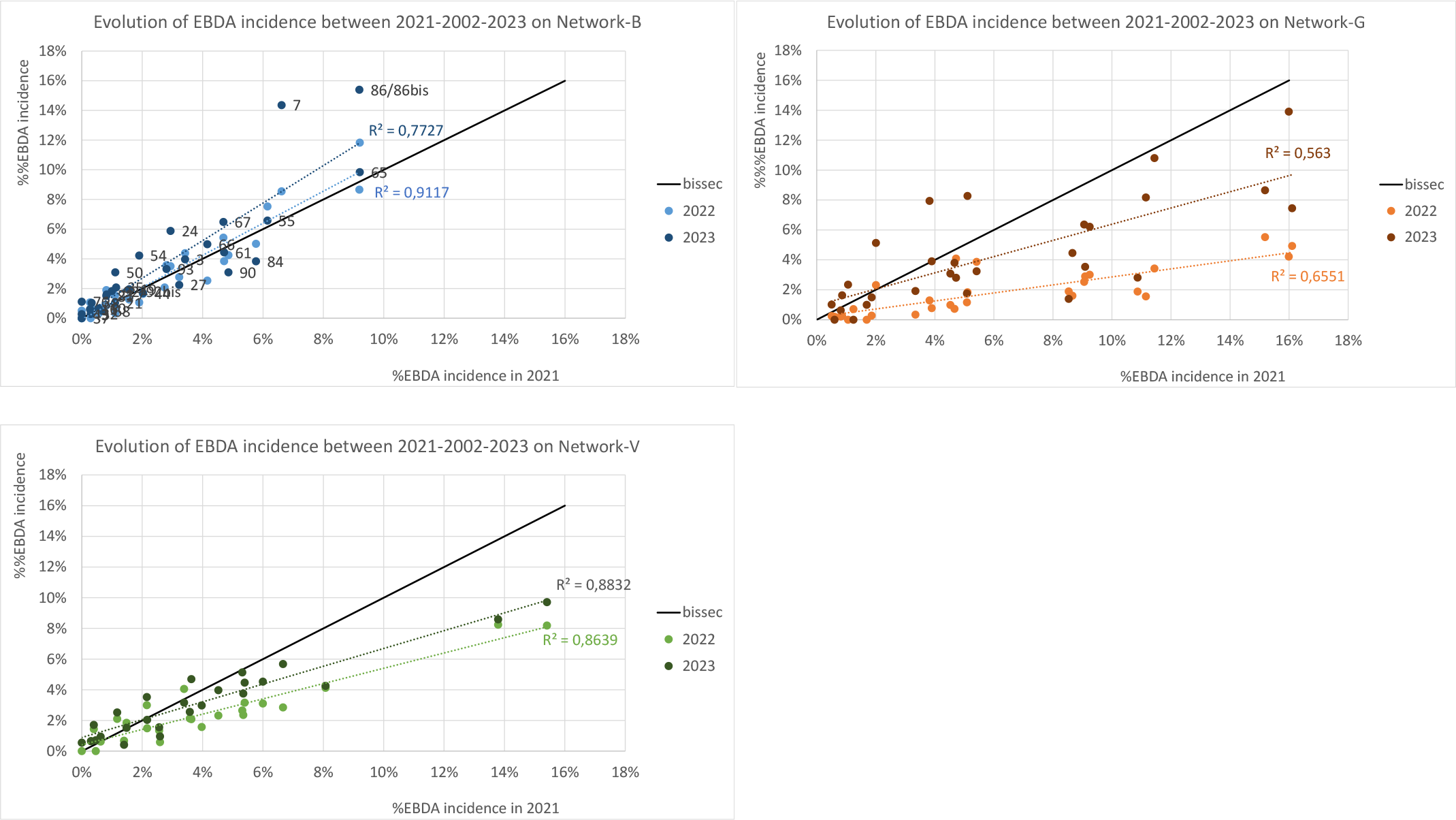
Comparison of total EBDA (including apoplexy) incidence between 2021 and 2022 / 2023 for each local plots Network-B, Network-V and Network-G. Black line curve figures the bisector.

#### 1.2. Climate in 2021, 2022 and 2023 on the 3 networks

Figure 3 shows the monthly average temperature and sum of rainfall for the vegetative period of April to August in 2021, 2022 and 2023.

**Figure 3.**
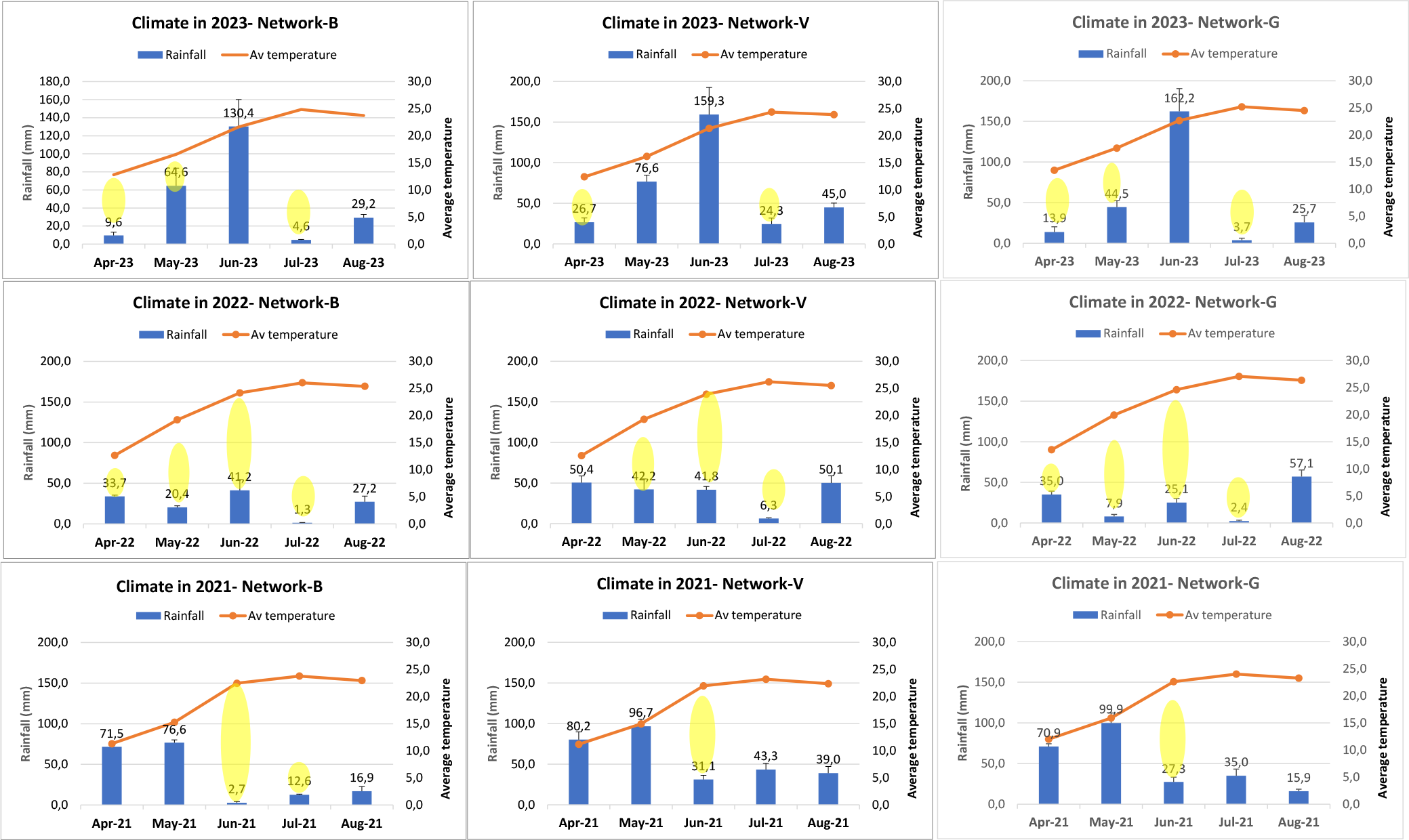
Monthly rainfall and average temperature over the 3 year- for each local plots Network-B, Network-V and Network-G. Yellow area is proposed as a standard to help compare and point out the differences between networks and years.

The same yearly tendencies are visible for all 3 networks with some differences between networks: 2021 was well watered in April and May but with a water shortage in June and July, more markedly visible in Network-B. 2022 was characterized by spring and summer drought, particularly strong in Network-G with a total of 60mm for the 4-month-period of April to July, when Network-B and Network-V received about 100 and 140mm of rainfall respectively. This corresponds to one half to one third of the 2021 rainfall amount on the same period. 2022 average temperatures (and evapotranspiration, *not shown*) are also higher. 2023 was characterized by a return of rainfall to higher amounts but with a discontinuous regime between a dry winter and April (*not shown*) followed by comfortable rainfall in May and June. The wet period of May and June 2023 was also characterized by stormy events that may have induced variability inside small geographical zones like the ones of our networks. This is shown on suppl. figure2. Little variability was visible inside the networks in 2021 and 2022, but in 2023, rainfall amounts between April and August in networks Network-G and Network-V could differ from 80 to 100 mm between extreme plots. According to Larignon (2020), these differences in rainfall amounts could be substantial enough to change the category of EBDA potential expression, especially in Network-V. For this reason, these differences were considered and integrated in the WB indexes calculation for 2023.

#### 1.3. Agronomical characterization of the networks and inter-relations between vigour and water, N and ground cover variables

A PCA was done to represent the inter-relations between the variables monitored in the study for the 3 networks and 2 years 2022 and 2023 taken together. Figure 4 (a) shows the inter-relations among the variables, with a colour per type of variable to help the reading (water, nitrogen, ground cover, production and vegetation scores at harvest and vigour) and figure 4 (b) shows a projection of the 3 networks on main F1 and F2 axis. Axis 1 accounts for 38% and the variance and both axis 1 and 2 almost 50%. Most of the indicators are correlated with axis 1 depicting water-N-ground cover relationships as well as grapevine yield and vegetation scores. Axis 2 is correlated with water status score at harvest, δC13 in 2022, as well as proportion of green cover in total ground cover in 2022, which can be interpreted as late water status information axis.

**Figure 4.**
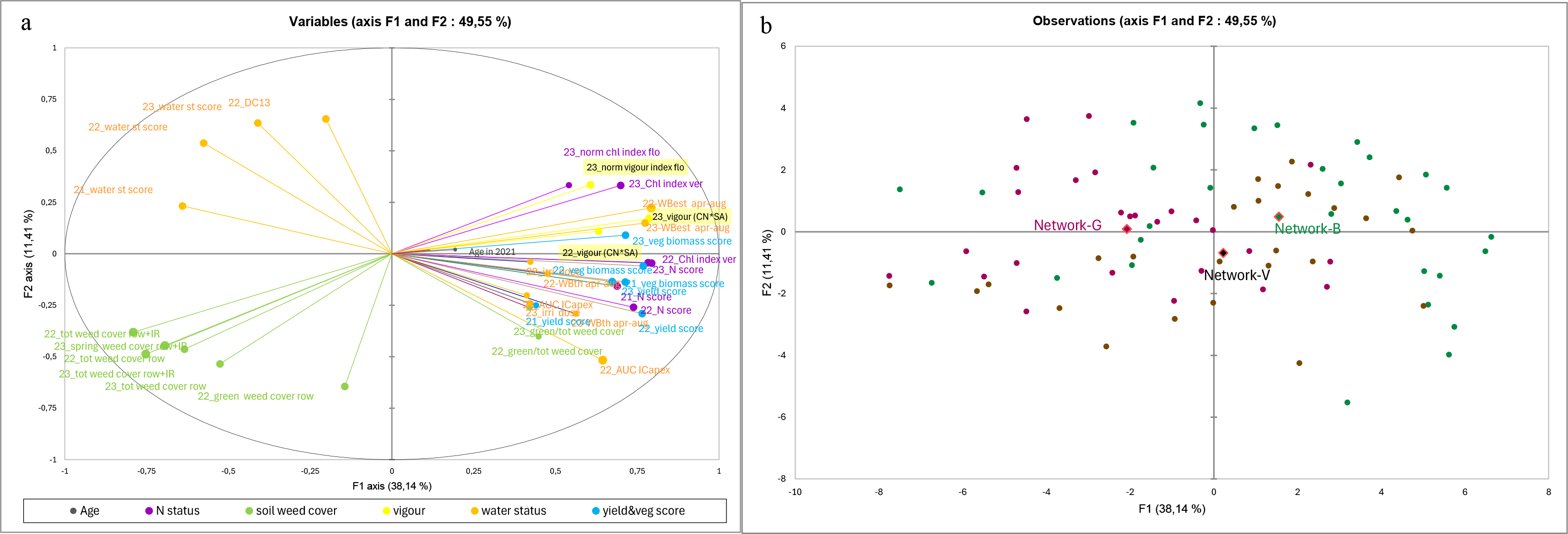
PCA performed on the matrix of the indicators describing water and nitrogen status, ground cover, production and vegetation score at harvest and vigour: (a) inter-relations between variables on the 2 first axis and (b) representation of the observations of the plots coloured by network type (Network-B in green, Network-G in purple or Network-V in brown) and the centroid of each network group of plots. On PCA of the variables: size of the circular label plot is proportional to the contribution of the variable on the axis it is best projected on. *Colour label legend for site descriptors categories: climate and water status variables in orange; grapevine nitrogen status in purple; weed cover in green; grapevine yield and vegetation scores at harvest in blue, vigour proxys ‘CN*sect’ in yellow; grapevine age in grey.

Vigour proxy CN*SA is strongly correlated with axis 1 and with ground cover and nitrogen status variables, as well as estimated water balance (WBest, itself intrinsically highly influenced by ground cover). On axis 1, vigour is also highly correlated to grapevine production and vegetation scores at harvest. Less water status indicators are correlated to axis 1, or with less strength.

The peculiarities of each network are visible on the PCA of observations (figure 4 b): Network-B and Network-G are significantly associated with axis 1. Network-B is located on the right side of axis 1, opposite to Network-G on the left side, Network-V being intermediate. Network-B is indeed characterized by more vigorous plots, less total ground cover on the row + inter-row, greater production in 2022. Network-G on the contrary has significantly more total ground cover (and less green cover), poorer nitrogen status for all 3 years and to a lesser extent, more deprived water status. Network-V is intermediate on most of the descriptors (ground cover, nitrogen, vigour) but has significantly higher water balances in 2022 and 2023 and also higher %green ground cover in 2022 and 2023 than the other 2 networks, the 2 latter probably due to the slightly higher rainfall amounts on this network those 2 years.

### 2. Relationships between vigour proxy, other plots descriptors and EBDA expression in 2022 and 2023

#### 2.1. Correlation matrix: best drivers for EBDA expression

Table 2 represents the correlations between the different descriptors and EBDA expression in 2022 (a) and 2023 (b) for all networks together and per network. As all the monitored variables are more or less correlated to one another (see fig 4 a), the p value (and rank of the p-value) was specified in order to represent the strength of the correlation. Last column recalls the indicators that are significantly correlated with vigour or not.

From a network to another, the number and nature of the variables significantly correlated to total EBDA expression vary, but: - “Vigour” proxy “CN*SA” is positively correlated and with a top rank value in 6 cases out of 8; in Network-V in 2022 and Network-G in 2023, there is only a tendency along with a p-value of 0.18;On all cases except Network-G in 2023, variables that are most correlated with %EBDA expression are also the ones that are most correlated with “vigour”, and with the same consistent trend (see Network-B in 2022 and 2023, Network-V in 2023 where almost all variables correlated to %EBDA display a cross in the column dedicated to correlation with vigour proxy). We can notice that some variables are recurrent: yield or vegetation biomass scores and ground cover indicators are present in every situation (global or per network). We can also notice that correlations with water status indicators are less present, except for estimated water balance (but this latter is heavily influenced by the level of ground vegetation cover).

- In Network-G in 2023, the strongest correlated variables are slightly different: these are indicators related with previous year vine water status. Indeed, AUC and δC13 in 2022 are correlated with 2023 %EBDA expression in a negative and positive way respectively. Production in 2022 is also correlated, but not with the positive direction it had when associated to vigour (like for Network-B and Network- V in 2023), but more likely reflecting 2022 water stress. Grapevine vigour still has a positive influence on EBDA expression on that network in 2023, visible on the positive correlation with the chlorophyll index at flowering 2023 and with CN*SA in 2022 and 2023 (in 2023 only in tendency, p=0.18). Nevertheless, it seems to be rather less influent than the correlation with previous year water stress variables. Note that water status in 2022 and vigour in 2023 are only partially correlated (2022 AUC and CN*SA 2023 positively correlated with p=0.05, 2022 AUC and 23_chl ind flo not correlated, p=0.57, *not shown*) and anyway, not showing the same direction effect on EBDA.

#### 2.2. Focus on relationship with vigour proxy CN*SA

Figure 5 specifies the relationship between vigour proxy “CN*SA” and EBDA incidence. We can see that the curves recurrently show the same type of shape, with a decreasing variance among EBDA incidence rates as grapevine vigour decreases: Network-B and Network-V in 2023 best account for that, where Network-G does not.

**Figure 5.**
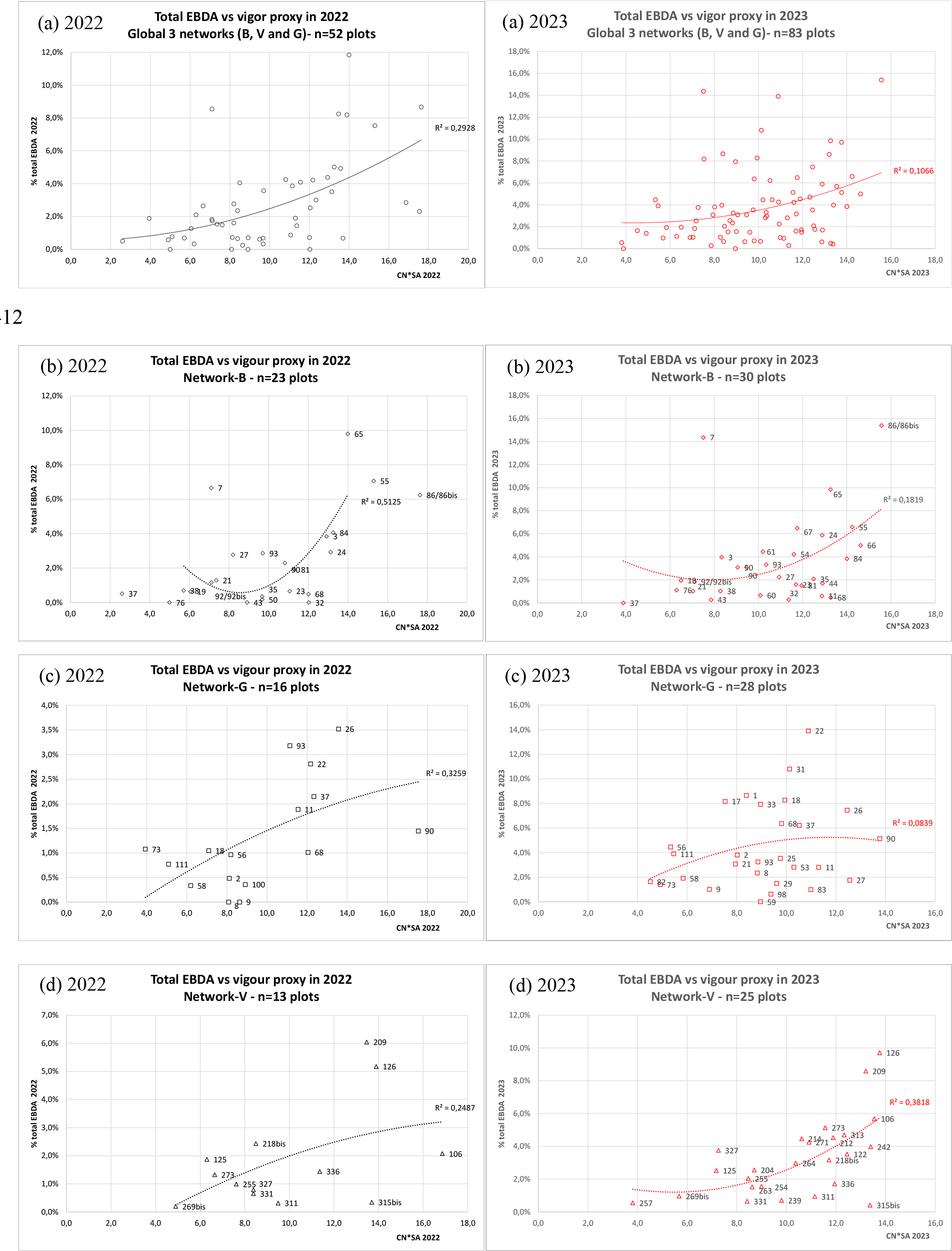
Bivariate relationships between vigour proxy CN*SA and total EBDA expression in 2022 (black) and 2023 (red) for all plots together (a) and for each local Network-B (b), Network-V (c) and Network-G (d) separately. Polynomial degree 2 trend curve (dotted-line) and R² determination coefficient. Labels= vineyard plot identification number.

This shape of curve can be read as (1) under a “CN*SA” proxy score of approximately 8, EBDA is low to even absent. (2) Rising from 8, the area described by the curve enlarges as vigour increases in such a way that for higher values of vigour, EBDA can be either high or not. For example, for Network-B and “CN*SA” scoring over 13, plots 65 and 86 show high level of EBDA in 2023, whereas plots 11, 44 and 68 do not. In Network-V, plots 125 and 209 show high level of EBDA in 2023 whereas plot 315bis does not.

Very low values of vigour are associated with no (or very little) esca expression, and this is visible on plots coming from all 3 networks: plots 37, 76, 19 or 21 in Network-B, plots 82 and 73 in Network- G, plots 269bis and 257 in Network-V. These plots all have in common to display high level of ground vegetation cover that is generally long-term permanent cover. Agronomically, these plots show low N status, little vegetative biomass, and very low yield. Plot 7 of Network-B seems to be an odd value, but not an outlier: despite a moderate vigour, EBDA expression in all 3 years is high. This would be worth investigating why.

As both (1) low vigour is associated with little EBDA symptom and (2) higher rates of symptoms are only seen for higher values of vigour (except for 2023 Network-G), we can conclude that vigour is associated to EBDA expression. But the fact that higher values of vigour can also be associated to low values of EBDA shows that other factor(s) are also influential on EBDA expression.

### 2.3. The case of Network-G in 2023

According to Table 2, the case of Network-G in 2023 seems to be slightly different from the others: the best correlation is shown with water status variables the year before, and in an opposite way from the one it would have had with vigour; nevertheless, a correlation with vigour is still visible in 2023. This incited us to try to analyse the EBDA expression using an interaction of the two factors.

**Table 1.** Variables and indicators monitored from 2021 through 2023 and their assessment periods.

| *Phenological period* | *Pre-flowering* | *Berry growth* | *Veraison* | *Harvest* | *Winter* |
| --- | --- | --- | --- | --- | --- |
| Climate data (temperature, rainfall, potential evapotranspiration) | 2021-2022-2023 |  |  |  |  |
| Water status: apex index | 2023 | 2022-2023 | 2022-2023 | 2022-2023 |  |
| Water status: $\delta C_{13}$ | | | | 2022* | |
| Nitrogen status: chlorophyll index | 2023 |  | 2022 |  |  |
| Ground vegetation cover: (total vs living, row vs inter-row) visual evaluation | 2023 | 2022-23 | 2022-23 | 2022-23 |  |
| Grapevine yield/ Vegetative biomass / water status / nitrogen status at harvest: visual evaluation |  |  |  | 2021-2022-2023 |  |
| EBDA / Apoplexy incidence |  |  |  | 2021-2022-2023 |  |
| Vine vigour: pruning weight proxy (CN*SA) |  |  |  |  | 2022*-2023 |

\* Only on half of the vineyard plots.

**Table 2.** Spearman correlation index, p-value and rank of the p-value between the different monitored indicators (colour label per type of variable, see below) and EBDA symptom expression (total EBDA including apoplexy form) for 2022 (a) and 2023 (b) successively for all plots together (“all 2022” and “all 2023”) or inside each network. Last column features a cross (“x”) for the variables that are significantly correlated with CN*SA vigour proxy (spearman correlation). Values in bold type= p value<0.10

| (a)<br>Variables | All 2022 |  |  |  |  | Network-B 2022 |  |  |  |  | Network-G 2022 |  |  |  |  | Network-V 2022 |  |  |  |
| --- | --- | --- | --- | --- | --- | --- | --- | --- | --- | --- | --- | --- | --- | --- | --- | --- | --- | --- | --- |
|  | Spearman corr index with 22_EBDA+apop | pvalue | p value-rank | variables correlated to 22_CN*SA |  | Spearman corr index with 22_EBDA+apop | pvalue | p value-rank | variables correlated to 22_CN*SA |  | Spearman corr index with 22_EBDA+apop | pvalue | p value-rank | variables correlated to 22_CN*SA |  | Spearman corr index with 22_EBDA+apop | pvalue | p value-rank | variables correlated to 22_CN*SA |
| Vine age in 2021 | -0,10 | 0,37 |  |  |  | **-0,33** | **0,07** | **10** |  |  | 0,15 | 0,46 |  |  |  | 0,06 | 0,76 |  |  |
| 21_yield score | 0,17 | 0,14 |  |  |  | 0,22 | 0,23 |  |  |  | 0,07 | 0,76 |  |  |  | 0,28 | 0,17 |  |  |
| 21_veg biomass score | **0,25** | **0,02** | **13** | **x** |  | **0,41** | **0,02** | **5** | **x** |  | 0,18 | 0,37 |  |  |  | 0,05 | 0,81 |  |  |
| 22_yield score | **0,27** | **0,01** | **11** | **x** |  | 0,29 | 0,12 |  |  |  | 0,06 | 0,75 |  |  |  | 0,33 | 0,10 |  |  |
| 22_veg biomass score | **0,41** | **0,00** | **4** | **x** |  | **0,41** | **0,02** | **6** | **x** |  | **0,34** | **0,09** | **3** | **x** |  | **0,38** | **0,06** | **6** | **x** |
| 21_water status score | **-0,19** | **0,09** | **15** |  |  | -0,18 | 0,32 |  |  |  | -0,05 | 0,80 |  |  |  | -0,10 | 0,63 |  |  |
| 22_Irrigation dose | 0,12 | 0,28 |  |  |  | 0,27 | 0,14 |  |  |  | -0,19 | 0,34 |  |  |  |  |  |  |  |
| 22_WBth apr-aug | **0,22** | **0,05** | **14** | **x** |  | **0,31** | **0,09** | **11** |  |  |  |  |  |  |  |  |  |  |  |
| 22_WBest apr-aug | **0,36** | **0,00** | **5** | **x** |  | **0,44** | **0,01** | **4** | **x** |  | 0,17 | 0,40 |  |  |  | **0,46** | **0,02** | **2** | **x** |
| 22_AUC | **0,33** | **0,00** | **7** | **x** |  | **0,39** | **0,03** | **8** |  |  | -0,11 | 0,61 |  |  |  | **0,45** | **0,02** | **4** |  |
| 22_DC13 | -0,05 | 0,72 |  |  |  | -0,30 | 0,24 |  |  |  | 0,20 | 0,44 |  |  |  | 0,14 | 0,60 |  |  |
| 22_water status score | **-0,25** | **0,02** | **12** | **x** |  | -0,28 | 0,13 |  |  |  | -0,09 | 0,64 |  |  |  | -0,15 | 0,47 |  |  |
| 21_N score | **0,41** | **0,00** | **3** |  |  | **0,60** | **0,00** | **2** | **x** |  | 0,14 | 0,48 |  |  |  | 0,32 | 0,12 |  |  |
| 22_chl index ver | **0,32** | **0,00** | **8** | **x** |  | **0,63** | **0,00** | **1** | **x** |  | -0,06 | 0,76 |  |  |  | 0,25 | 0,23 |  |  |
| 22_N score | **0,31** | **0,01** | **9** | **x** |  | 0,31 | 0,11 |  |  |  | 0,19 | 0,36 |  |  |  | **0,39** | **0,06** | **5** |  |
| 22_tot weed cover row | **-0,44** | **0,00** | **1** | **x** |  | **-0,40** | **0,03** | **7** | **x** |  | -0,31 | 0,13 |  |  |  | **-0,47** | **0,02** | **1** | **x** |
| 22_green weed cover row | **-0,27** | **0,01** | **10** | **x** |  | -0,13 | 0,47 |  |  |  | **-0,38** | **0,06** | **2** |  |  | **-0,46** | **0,02** | **3** |  |
| 22_tot weed cover row+IR | **-0,34** | **0,00** | **6** | **x** |  | **-0,36** | **0,05** | **9** | **x** |  | -0,18 | 0,38 |  |  |  | **-0,46** | **0,02** | **3** | **x** |
| 22_green/tot weed cover | 0,03 | 0,80 |  |  |  | 0,18 | 0,51 |  |  |  | -0,13 | 0,52 |  |  |  | 0,02 | 0,93 |  |  |
| 22_CN*SA | **0,53** | **0,00** | **2** |  |  | **0,55** | **0,01** | **3** |  |  | **0,65** | **0,01** | **1** |  |  | 0,40 | 0,18 |  |  |
| 22_EBDA+apop | **1,00** | **0,00** |  |  |  | **1,00** | **0,00** |  |  |  | **1,00** | **0,00** |  |  |  | **1,00** | **0,00** |  |  |

| (b)<br>Variables | All 2023 |  |  |  | Network-B 2023 |  |  |  | Network-G 2023 |  |  |  | Network-V 2023 |  |  |  |
| --- | --- | --- | --- | --- | --- | --- | --- | --- | --- | --- | --- | --- | --- | --- | --- | --- |
|  | Spearman corr<br>index with<br>23_EBDA+apop | pvalue | pvalue-<br>rank | variables<br>correlated to<br>23_CN*SA | Spearman corr<br>index with<br>23_EBDA+apop | pvalue | pvalue-<br>rank | variables<br>correlated to<br>23_CN*SA | Spearman corr<br>index with<br>23_EBDA+apop | pvalue | pvalue-<br>rank | variables<br>correlated to<br>23_CN*SA | Spearman corr<br>index with<br>23_EBDA+apop | pvalue | pvalue-<br>rank | variables<br>correlated to<br>23_CN*SA |
| Vine age in 2021 | -0,18 | 0,11 |  |  | -0,37 | 0,05 | 8 |  | -0,14 | 0,51 |  |  | 0,06 | 0,79 |  |  |
| 21_yield score | 0,09 | 0,42 |  |  | 0,31 | 0,10 | 14 | x | -0,21 | 0,32 |  |  | 0,07 | 0,72 |  |  |
| 21_veg biomass score | 0,09 | 0,41 |  |  | 0,29 | 0,12 |  |  | 0,02 | 0,94 |  |  | 0,03 | 0,87 |  |  |
| 22_yield score | 0,03 | 0,80 |  |  | 0,20 | 0,29 |  |  | -0,41 | 0,04 | 3 | x | 0,37 | 0,07 | 6 | x |
| 22_veg biomass score | 0,22 | 0,05 | 7 | x | 0,36 | 0,05 | 9 | x | 0,13 | 0,53 |  |  | 0,27 | 0,20 |  |  |
| 23_yield score | 0,21 | 0,06 | 8 | x | 0,32 | 0,08 | 13 | x | -0,16 | 0,45 |  |  | 0,42 | 0,04 | 3 | x |
| 23_veg biomass score | 0,12 | 0,28 |  |  | 0,12 | 0,54 |  |  | 0,07 | 0,75 |  |  | 0,18 | 0,41 |  |  |
| 21_water status score | 0,07 | 0,55 |  |  | -0,06 | 0,76 |  |  | 0,13 | 0,55 |  |  | -0,09 | 0,67 |  |  |
| 22_Irrigation dose | -0,06 | 0,58 |  |  | 0,14 | 0,45 |  |  | -0,32 | 0,11 |  |  |  |  |  |  |
| 22_WBth apr-aug | -0,12 | 0,29 |  |  | 0,14 | 0,47 |  |  |  |  |  |  |  |  |  |  |
| 22_WBest apr-aug | 0,07 | 0,54 |  |  | 0,27 | 0,15 |  |  | 0,09 | 0,66 |  |  | 0,35 | 0,09 | 9 | x |
| 22_AUC | 0,03 | 0,78 |  |  | 0,20 | 0,28 |  |  | -0,45 | 0,03 | 2 | (x) | 0,31 | 0,13 |  |  |
| 22_DC13 | 0,26 | 0,06 | 9 |  | 0,00 | 1,00 |  |  | 0,59 | 0,02 | 1 |  | 0,12 | 0,65 |  |  |
| 22_water status score | 0,02 | 0,89 |  |  | -0,07 | 0,70 |  |  | 0,30 | 0,15 |  |  | -0,19 | 0,35 |  |  |
| 23_Irrigation dose | -0,14 | 0,21 |  |  | -0,07 | 0,72 |  |  | -0,32 | 0,12 |  |  |  |  |  |  |
| 23_WBth apr-aug | -0,05 | 0,66 |  |  | 0,10 | 0,62 |  |  | 0,23 | 0,26 |  |  | 0,10 | 0,63 |  |  |
| 23_WBest apr-aug | 0,05 | 0,66 |  |  | 0,15 | 0,44 |  |  | 0,27 | 0,19 |  |  | 0,22 | 0,30 |  |  |
| 23_AUC | 0,13 | 0,25 |  |  | 0,19 | 0,33 |  |  | -0,05 | 0,80 |  |  | 0,12 | 0,56 |  |  |
| 23_water status score | 0,07 | 0,55 |  |  | -0,07 | 0,69 |  |  | 0,29 | 0,16 |  |  | -0,01 | 0,94 |  |  |
| 21_N score | 0,20 | 0,07 | 11 | x | 0,38 | 0,04 | 7 | x | 0,07 | 0,76 |  |  | 0,24 | 0,25 |  |  |
| 22_chl index ver | 0,02 | 0,85 |  |  | 0,51 | 0,00 | 2 | x | -0,30 | 0,14 |  |  | 0,23 | 0,28 |  |  |
| 22_N score | 0,06 | 0,62 |  |  | -0,01 | 0,97 |  |  | -0,05 | 0,81 |  |  | 0,29 | 0,16 |  |  |
| 23_chl index flo | 0,04 | 0,73 |  |  | 0,21 | 0,26 |  |  | 0,38 | 0,07 | 5 | x | 0,33 | 0,15 |  |  |
| 23_chl index ver | 0,03 | 0,78 |  |  | 0,08 | 0,66 |  |  | 0,14 | 0,51 |  |  | 0,07 | 0,75 |  |  |
| 23_N score | 0,17 | 0,13 |  |  | 0,40 | 0,03 | 5 | x | -0,26 | 0,21 |  |  | 0,36 | 0,08 | 7 | x |
| 22_tot weed cover row | -0,28 | 0,01 | 4 | x | -0,39 | 0,03 | 6 | x | -0,18 | 0,38 |  |  | -0,38 | 0,06 | 5 | x |
| 22_green weed cover row | -0,26 | 0,02 | 6 | x | -0,35 | 0,06 | 10 |  | -0,28 | 0,17 |  |  | -0,46 | 0,02 | 2 | x |
| 22_tot weed cover row+IR | -0,12 | 0,27 |  |  | -0,34 | 0,07 | 12 | x | -0,10 | 0,64 |  |  | -0,35 | 0,09 | 8 | x |
| 22_green/tot weed cover | -0,23 | 0,07 | 10 | x | -0,14 | 0,63 |  |  | -0,22 | 0,29 |  |  | -0,14 | 0,51 |  |  |
| 23_tot weed cover row | -0,17 | 0,15 |  |  | -0,20 | 0,30 |  |  | -0,31 | 0,13 |  |  | -0,34 | 0,13 |  |  |
| 23_green weed cover row | -0,26 | 0,02 | 5 | x | -0,35 | 0,06 | 11 | x | -0,15 | 0,47 |  |  | -0,40 | 0,05 | 4 | x |
| 23_tot weed cover row+IR | -0,17 | 0,13 |  |  | -0,30 | 0,11 |  |  | -0,27 | 0,19 |  |  | -0,28 | 0,17 |  |  |
| 23_green/tot weed cover | -0,09 | 0,42 |  |  | 0,10 | 0,60 |  |  | -0,26 | 0,21 |  |  | 0,29 | 0,16 |  |  |
| 22_CN*SA | 0,45 | 0,00 | 1 | x | 0,54 | 0,01 | 3 | x | 0,53 | 0,05 | 4 | x | 0,26 | 0,39 |  |  |
| 23_CN*SA | 0,33 | 0,00 | 3 |  | 0,41 | 0,02 | 4 |  | 0,28 | 0,18 |  |  | 0,57 | 0,00 | 1 |  |
| 23_EBDA+apop | 1,00 | 0,00 |  |  | 1,00 | 0,00 |  |  | 1,00 | 0,00 |  |  | 1,00 | 0,00 |  |  |

Figure 6 presents the bivariate correlation curves between EBDA expression in 2023 in Network-G and two indicators related to vigour in 2023 (chlorophyll index at flowering and CN*SA, fig.4 a) and to water stress in 2022 (AUC and δC13, fig.4 b).

**Figure 6.**
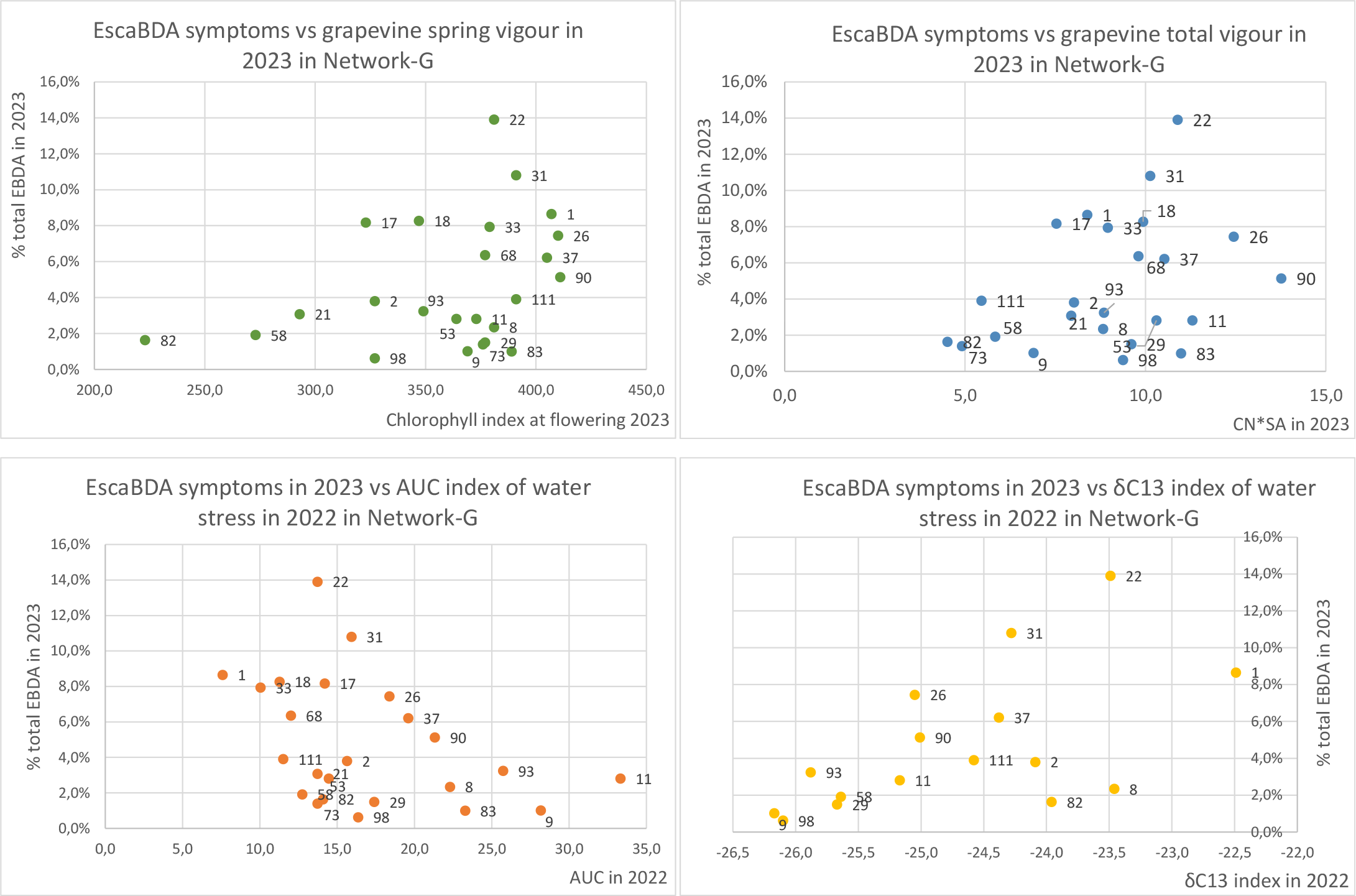
Bivariate relationships between EBDA expression in 2023 in Network-G and 2 indicators related to vigour in 2023 (chlorophyll index at flowering and CN*SA, a) and to water status in 2022 (AUC and δC13, b). Labels= vineyard plot identification number.

We can visualize on these graphs the positive effect of the current year vigour and the positive effect of the water stress in the year before on the incidence of EBDA symptoms in 2023 in this network. Here again, the same type of curve shape is noticeable for both variables (vigour and water stress) indicating that apart from those 2 variables, there are still some additional hidden factor(s) acting on EBDA expression.

The analyse of the respective position of the vineyard plots on either the vigour or the water status curves shows that there is a decoupling between vigour and water status indicators showing distinct combinations depending on the vineyard plot:

- Some plots that showed rather high vigour in 2023, especially in spring, may have suffered either low water stress in 2022 (see plots 26-37-90, EBDA expression is 5 to 8% in 2023) or high water stress in 2022 (plots 31, 22 and 1, EBDA 8 to14% in 2023);

- On the contrary, plots with moderate vigour (especially in spring) in 2023 may have suffered water stress (plots 17-18, EBDA 8%) or not (like irrigated plots 8-9-11, EBDA 1 to 3% or 93, 2 and 21 plots, EBDA 3-4%) in 2022.

Worst situation for EBDA expression seems to be found when the plot has suffered high water stress in 2022 and still shows substantial vigour in 2023. This is the case of plots 1, 22 and 33 with high water stress in 2022 but good spring vigour in 2023, showing high levels of EBDA expression in 2023 (8-14%). We can notice that plot 22 also showed an elevated total vigour (CN*SA in 23), whereas plots 1 and 33 rather moderate ones, suggesting that active growth on plot 22 lasted longer in 2023; to what extent this may be related to the highest disease expression on plot 22 would need to be confirmed. Those 3 plots have in common to be located on the site of a former pine forest with clayey soils that are probably fertile in springtime (due to high levels of organic matter) but dry out quickly in the summer.

Plots 90 and 26 are very fertile. They had no water stress in 2022 and had high EBDA expression in 2023 (6-8%), but still a step below the plots that experienced water stress in 2022.

We can notice that, as long as the vigour is very low both in the spring and for total vigour (plots 58, 82), EBDA incidence rates seem to be low as well (2%). Plots 21 and 111 show low vigour only on one of the two indicators (21 in the spring, 111 in total vigour), and slightly higher levels (3-4%), though still moderate, of EBDA expression in 2023.

We can still visualize the action of the ‘unknown factor(s)’ on plots 98, 2 and 17 for instance, that showed similar moderate vigour in 2023 with similar low to moderate water stress in 2022 but still displayed distinct levels of EBDA in 2023 (<1%, 4% and 8% respectively). The case of plot 83 is also interesting because it is a rather vigorous plot with no water stress in 2022, and very low EBDA expression.

## Discussion

The cases from the three networks studied over two contrasting years offer a diverse range of viticultural practices (such as irrigation, permanent ground vegetation cover and production goals) as well as environmental conditions, in particular climate: the 2022 season was dry, while 2023 showed a dry spring followed by rainy May and June. These were conducive conditions to test the “vigour” hypothesis in different contexts.

This study shows that grapevine “vigour” was correlated to EBDA expression in 2022 and 2023 in 5 out of 6 situations, with a curve type shape meaning that:

- When the vigour is very low, there is only little to no EBDA expression. These are plots with particularly low vigour (and production), generally due to long term prior co-existence of vine and weeds over the entire soil surface (inter-row and row). When the vigour increases, the expression of EBDA increases too but to a variable extent, so that it is possible to find vigorous plots with either high or low EBDA expression. However, in these situations, the highest rates of EBDA incidence are always associated with high vigour.

- In our study, the best indicator of vigour is the total cane biomass proxy CN*SA; but this indicator itself is recurrently correlated with vine nitrogen status, ground cover, and grapevine production and vegetation scores. These indicators are also correlated to EBDA expression, supporting the central role of “vigour” in EBDA expression. Vine water status indicators show weaker correlations.

- The results suggest that vigour is not the only variable implicated in EBDA expression, as expected from these complex multi-factorial diseases. Additional variables are needed to account for vigorous and EBDA symptom-less plots. These variables must be strong enough to counteract the effect of the vigour. They also might act priorly to vigour and be necessary (still not sufficient) to lead to high level of symptom expression. Winter pruning could be one of those influential variables, affecting the internal health of the wood relatively to subsequent desiccation and necrosis amounts. For example, in Network-G, plots 83 and 98, that show particularly low EBDA expression relative to their vigour and water stress levels, use training systems respectful of sap flow pathways (respectively goblet and cordon with elongated branches). Assessing pruning quality in plots with similar vigour but different EBDA expression levels could provide valuable insights.

- In one case out of six, the one of Network-G in 2023, vigour (and related variables) was not the primary factor driving EBDA expression. The main driver seemed to be water stress in the previous year, suggesting that the water stress in 2022 positively influenced EBDA expression in 2023. Vigour was still influential but played a lesser role. Besides, the vigour indicator best correlated was chlorophyll index at flowering, which may suggest an early influence of grapevine vigour in the season. In this network, the plots that displayed the highest levels of symptoms in 2023 were the ones that suffered the highest water stress in 2022 but still showed rather high levels of spring vigour in 2023. This potential dual action would need further confirmation with additional years and locations.

It is interesting to note that it is Network-G, characterized by the highest permanent ground vegetation cover percentage, was affected by this water stress type of influence. In 2022, we observed in distinct south-eastern mediterranean situations that the vineyards that first began to suffer from water stress in early June were the ones showing substantial ground vegetation cover. Consequently, it is plausible that high weed cover significantly burdened grapevine physiology in 2022 in our Mediterranean region. Besides, Network-G had the highest frequency of plots affected by fanleaf symptoms, a debilitating disease caused by *Nepoviruses* propagated by soil nematodes, leading to discoloration, abnormalities, and stunted vegetation (*not shown*). To what extent this additional charge for vines can be involved in EBDA over-expression would need to be investigated. In Network-B, plot n°7 had an outlying behaviour as it showed rather moderate vigour compared to its high level of disease expression. Yet, it is a plot with all inter-rows covered with permanent grass and no irrigation. To what extent it is possible to think that this vineyard is set in a similar situation as the ones of Network- G would also be worth investigating.

These results of the positive effect of vigour on EBDA symptoms are in accordance with Dumot (2022) in the French Cognac region, and Gastou et al. (2024) that in addition, highlighted the same kind of relationship with both pruning weight and water use efficiency. Monod et al. (2023) observed a positive relationship between EBDA incidence and soil water holding capacity, which is consistent with a positive influence of ’vigour.’ However, in this study, the correlation with the pruning weight vigour proxy itself was weaker. This suggests that, in those three cases, there is a trend to more EBDA incidence when the ‘vigour’ is higher, but depending on the year, the geographical, climate and viticultural situation, the best proxies differ. It may also suggest intricated mechanisms between soil capacity, water / N uptake and vigour that may depend on the peculiarities of each viticultural region (i.e. climate, soil, vine cultivars, and rootstocks).

The preliminary observation made in Network-G in 2023, outlining a correlation between EBDA expression and occurrence of water stress the year before, is also consistent with testimonies collected from different French regions like Alsace, Jura or Beaujolais vineyards (*personal communication*). Interestingly, permanent ground vegetation cover is frequent in these vineyards, as it is in Network- G.

Duplicating similar monitoring across a broader range of viticultural situations, including years, regions and cultivar/rootstock genotypes would be interesting to confirm the two putative profiles observed in our study.

In this study, two influent factors proved to be involved in EBDA expression: vine vigour on one hand, and previous year water stress on the other. Other studies have highlighted additional favourable factors, such as harvest date (Kuntzmann et al., 2013), water stress (Calvo-Garrido et al., 2021) or on the contrary, water availability (Bortolami et al., 2021). This suggests that, beyond these variables, there might be a more upstream factor in relation to the global vine physiology that could be the real causal variable explaining EBDA symptom expression. To account for that, it is possible to think of grapevine carbon or nitrogen balance, and source-sink relationships: in that perspective, vigour, related to high vegetative biomass and grapes, is indeed a strong sink for C assimilates. Water stress the year before, on the other hand, can limit the sources production by preventing proper restorage of C. The apparent discrepancies between results from different studies as well as the best drivers differing between regions and years might originate from the fact that C balance in grapevine is multi-factorial (so generalizing the effect of one factor alone is not easy to do because of its interaction with the remaining factors) or because factors have nonlinear effects or else because of the variation of the initial status of the vine from one year to another. The link between C balance in grapevine and EBDA symptom expression can be assumed considering defence metabolites. These defence metabolites are often considered in plants to derive from C primary metabolism and as a result, source-sink relationships may be highly influent. They might as well rely on starch and other C reserves, particularly the ones stored in the trunk in the vicinity of wood inhabiting fungal agents of trunk diseases before or during the outburst of a symptomatic event. The allocation for defence metabolites might thus be favoured by an excess of C available, once the priority sinks have been supplied. As for the time period of susceptibility, springtime has proven to be an important stage: Spagnolo et al. (2014) showed that flowering is a particularly susceptible stage when contaminating green shoots with two *Botryosphaeriaceae* species. Besides, climate analysis has also proven that the temperature and rainfall conditions during the 2 month-period preceding the symptomatic events, i.e. in our conditions April to May, are critical (Fréjaville et al., 2022; Larignon, 2009). We can assume that springtime is a critical period that could be acting either on triggering or on predisposing the vine to the subsequent symptomatic event. This, along with the previous ones, are hypothesis that would be worth investigating in further studies.

In this study, we identified the role of grapevine vigour as an influential factor on EBDA and this opens new perspectives of applications for growers, like assessing whether vigour management could be a sustainable way to reduce trunk disease expression and mortality for the affected plots. But this raises questions: will vigour management induce a reaction on EBDA expression and if so, what is the expected time frame for observing a response? What vigour management techniques and driving should be employed (e.g. ground vegetation cover management, fertilization, irrigation …)? Is a trade-off with grapevine yield achievable, i.e. can we drive vigour reduction in a susceptible vineyard without too much yield inflexion relatively to the growers’ objectives? And what about the subsequent grape and wine typicity in the current climate change context?

Further work is encouraged to address these questions.

## Acknowledgements

The authors thank the vine growers for their helpful availability in this study, as well as the partner wineries for their collaboration.

The authors also thank Cédric Moisy, Jean-Yves Cahurel and Philippe Larignon for their kind and helpful proofreading as well as Sam Schrock for his patience and relevance in checking up the English.

This work was part of the Dep Grenache project from the French *Plan National de Lutte contre le Dépérissement du Vignoble (PNDV)* and was made possible thanks to French Ministry of Agriculture (CASDAR, FranceAgriMer) and CNIV funding. The French Ministry of Agriculture and FranceAgriMer cannot be held liable for any damages resulting from the use of the information contained in this article.

## References

1. Abidon, C., Malblanc, S., & Funfrock, N. (Novembre 2019). Bilan des maladies du bois en Alsace: 2019, mise en place d’un nouvel observatoire. Les Vins D’alsace, 11.

2. Bertsch, C., Ramírez-Suero, M., Magnin-Robert, M [M.], Larignon, P., Chong, J., Abou-Mansour, E., Spagnolo, A., Clément, C., & Fontaine, F. (2013). Grapevine trunk diseases: Complex and still poorly understood. Plant Pathology, 62(2), 243–265. 10.1111/j.1365-3059.2012.02674.x

3. Bortolami, G., Gambetta, G. A., Cassan, C., Dayer, S., Farolfi, E., Ferrer, N., Gibon, Y., Jolivet, J., Lecomte, P., & Delmas, C. E. L. (2021). Grapevines under drought do not express esca leaf symptoms. Proceedings of the National Academy of Sciences, 118(43). 10.1073/pnas.2112825118

4. Calvo-Garrido, C., Songy, A., Marmol, A., Roda, R., Clément, C., & Fontaine, F. (2021). Description of the relationship between trunk disease expression and meteorological conditions, irrigation and physiological response in Chardonnay grapevines. OENO One, 55(2), 97–113. 10.20870/oeno-one.2021.55.2.4548

5. Calzarano, F., Fabio, O., Baranek, M., & Di Marco, S. (2018). Rainfall and temperature influence expression of foliar symptoms of grapevine leaf stripe disease (esca complex) in vineyards. Phytopathologia Mediterranea, 57(3), 488–505. 10.14601/Phytopathol_Mediterr-23787

6. Carbonneau, A. (1994). Le zonage des potentialités viticoles à l’échelle de l’Union Européenne. Le Progrès Agricole Et Viticole, 111. https://agris.fao.org/search/en/providers/123819/records/647362a1e17b74d22253c394

7. Champagnol, F. (1984). Elements de physiologie de la vigne et de viticulture generale.

8. Claverie, M., Notaro, M., Fontaine, F., & Wery, J. (2020). Current knowledge on Grapevine Trunk Diseases with complex etiology: a systemic approach. Phytopathologia Mediterranea, 59(1), 29–53. 10.36253/phyto-11150

9. Dal, F., Bricaud, E., Chagnon, L., & Daulny, B. (2008). Relation entre qualite de la taille et deperissement des vignes. Exemple de l’Esca. Le Progrès Agricole Et Viticole, 125(22), 602–608.

10. Destrac-Irvine, A., Goutouly, J. P., Laveau, C., & Guérin Dubrana, L. (2007). L’écophysiologie de la vigne – Mieux comprendre les maladies de dépérissement. L’union Girondine Des Vins De Bordeaux, 1035, 28–32.

11. Dewasme, C., Mary, S., Darrieutort, G [Guillaume], Roby, J.-P., & Gambetta, G. A. (2022). Long-Term Esca Monitoring Reveals Disease Impacts on Fruit Yield and Wine Quality. Plant Disease, 106(12), 3076– 3082. 10.1094/PDIS-11-21-2454-RE

12. Dumot, V. (Mai 2022). Fertilisation azotée: les enseignements d’un essai de 20 ans. UGNIC L’avenir Du Cognac, 71.

13. Etienne, L., Fabre, F., Martinetti, D., Frank, E., Michel, L., Bonnardot, V., Guérin Dubrana, L., & Delmas, C. E. L. (2024). Exploring the role of cultivar, year and plot age in the incidence of grapevine trunk diseases: Insights from 20 years of regional surveys in France. BioRxiv, 2024.03.19.585220. 10.1101/2024.03.19.585220

14. Fernandez, R., Le Cunff, L., Mérigeaud, S., Verdeil, J.-L., Perry, J., Larignon, P., Spilmont, A.-S., Chatelet, P., Cardoso, M., Goze-Bac, C., & Moisy, C. (2024). End-to-end multimodal 3D imaging and machine learning workflow for non-destructive phenotyping of grapevine trunk internal structure. Scientific Reports, 14(1), 5033. 10.1038/s41598-024-55186-3

15. Fischer, M., & Kassemeyer, H. H. (2012). Water regime and its possible impact on expression of Esca symptoms in Vitis vinifera: growth characters and symptoms in the greenhouse after artificial infection with Phaeomoniella chlamydospora. VITIS-Journal of Grapevine Research, 51(3), 129.

16. Fischer, M., & Peighami-Ashnaei, S. (2019). Grapevine, esca complex, and environment: the disease triangle. Phytopathologia Mediterranea, 58(1), 17–37. 10.13128/Phyto-pathol_Mediterr-25086

17. Fontaine, F., Pinto, C., Vallet, J., Clément, C., Gomes, A. C., & Spagnolo, A. (2016). The effects of grapevine trunk diseases (GTDs) on vine physiology. European Journal of Plant Pathology, 144(4), 707–721. 10.1007/s10658-015-0770-0

18. Fréjaville, T., Guérin Dubrana, L., Larignon, P., Lecomte, P., & Delmas, C. E. (2022). Short-term relationships between climate and grapevine trunk diseases in southern French vineyards. In Terclim2022 (Chair), *Terclim2022*, Bordeaux. https://ives-openscience.eu/wp-content/uploads/2023/02/f18-flash-frejavillet_ok.pdf

19. Gastou, P., Destrac Irvine, A., Arcens, C., Courchinoux, E., This, P., van Leeuwen, C., & Delmas, C. E. L. (2024). Large gradient of susceptibility to esca disease revealed by long-term monitoring of 46 grapevine cultivars in a common garden vineyard. BioRxiv, 2024.03.11.584376. 10.1101/2024.03.11.584376

20. Gaudillère, J.-P., van Leeuwen, C., & Trégoat, O. (2001). The assessment of vine water uptake conditions by ^13^c/^12^c discrimination in grape sugar. OENO One, 35(4), 195. 10.20870/oeno-one.2001.35.4.984

21. Gramaje, D., Úrbez-Torres, J. R., & Sosnowski, M. R. (2018). Managing grapevine trunk diseases with respect to etiology and epidemiology: current strategies and future prospects. Plant Disease, 102(1), 12–39. 10.1094/PDIS-04-17-0512-FE

22. Grosman, J., & Doublet, B. (2012). Maladies du bois de la vigne: synthèse des dispositifs d’observation au vignoble, de l’observatoire 2003-2008 au réseau d’épidémiosurveillance actuel. Phytoma-La Défense Des Végétaux(651), 31–35.

23. Guérin Dubrana, L., Fontaine, F., & Mugnai, L. (2019). Grapevine trunk disease in European and Mediterranean vineyards: occurrence, distribution and associated disease-affecting cultural factors. Phytopathologia Mediterranea, 58(1), 49–71. 10.13128/Phytopathol_Mediterr-25153

24. Kraus, C., Rauch, C., Kalvelage, E. M., Behrens, F. H., d’Aguiar, D., Dubois, C., & Fischer, M. (2022). Minimal versus Intensive: How the Pruning Intensity Affects Occurrence of Grapevine Leaf Stripe Disease, Wood Integrity, and the Mycobiome in Grapevine Trunks. Journal of Fungi, 8(3), 247. 10.3390/jof8030247

25. Kuntzmann, P., Barbe, J., Maumy-Bertrand, M., & Bertrand, F. (2013). Late harvest as factor affecting esca and Botryosphaeria dieback prevalence of vineyards in the Alsace region of France. Vitis, 52(4),, 197– 204.

26. La Fuente, M. de, Fontaine, F., Gramaje, D., Armengol, J., Smart, R., Nagy, Z. A., Borgo, M., Rego, C., & Corio-Costet, M.-F. (2016). Grapevine trunk diseases. A review (1st edition). International Organisation of Vine and Wine (OIV).

27. Larignon, P. (2009). Y a-t-il un lien entre climat et expression du Black Dead Arm? Identification des facteurs climatiques favorisant l’expression des symptômes. Phytoma-La Défense Des Végétaux(628), 27–29.

28. Larignon, P. (2020). Impact du changement climatique sur l’expression des symptômes de l’esca/bda dans le vignoble français. Revue Des Oenologues Et Des Techniques Vitivinicoles Et Oenologiques, 47 (*176*), 26–29.

29. Larignon, P., Fontaine, F., Farine, S., Clément, C., & Bertsch, C. (2009). Esca et Black Dead Arm: deux acteurs majeurs des maladies du bois chez la Vigne. Comptes Rendus De L’académie Des Sciences- BIOLOGIES, 332, 765–783.

30. Lecomte, P., Bénétreau, C., Diarra, B., Meziani, Y., Delmas, C., & Fermaud, M. (2023). Logistic modeling of summer expression of esca symptoms in tolerant and susceptible cultivars in Bordeaux vineyards. OENO One, 58(1). 10.20870/oeno-one.2024.58.1.7571

31. Lecomte, P., Darrieutort, G [G.], Laveau, C., Louvet, G., Goutouly, J. P., Rey, P [P.], & Guérin Dubrana, L. (2011). Impact of biotic and abiotic factors on the development of esca decline disease. In *«Integrated Protection and Production in Viticulture»*.

32. Lecomte, P., Diarra, B., Carbonneau, A., Patrice, R. E., & Chevrier, C. (2019). Esca of grapevine and training practices in France: results of a 10-year survey. Phytopathologia Mediterranea, 57(3), 472–487. 10.14601/Phytopathol_Mediterr-22025

33. Li, S., Bonneu, F., Chadoeuf, J., Picart, D., Gégout-Petit, A., & Guérin Dubrana, L. (2017). Spatial and Temporal Pattern Analyses of Esca Grapevine Disease in Vineyards in France. Phytopathology, 107(1), 59–69. 10.1094/PHYTO-07-15-0154-R

34. Marchi, G., Peduto, F., Mugnai, L., Di Marco, S., Calzarano, F., & Surico, G. (2006). Some observations on the relationship of manifest and hidden esca to rainfall. Phytopathologia Mediterranea, 45(4), 117–126. 10.1400/52267

35. Mondello, V., Songy, A., Battiston, E., Pinto, C., Coppin, C., Trotel-Aziz, P., Clément, C., Mugnai, L., & Fontaine, F. (2018). Grapevine trunk diseases: a review of fifteen years of trials for their control with chemicals and biocontrol agents. Plant Disease, 102(7), 1189–1217. 10.1094/PDIS-08-17-1181-FE

36. Monod, V., Zufferey, V., Wilhelm, M., Viret, O., Gindro, K., Croll, D., & Hofstetter, V. (2023). A systemic approach allows to identify the pedoclimatic conditions most critical in the susceptibility of a grapevine cultivar to esca/Botryosphaeria dieback. BioRxiv, 2023.05.23.541976. 10.1101/2023.05.23.541976

37. Mugnai, L., Graniti, A., & Surico, G. (1999). Esca (black measles) and brown wood-streaking: two old and elusive diseases of grapevines. Plant Disease, 83(5), 404–418. 10.1094/PDIS.1999.83.5.404

38. Pichon, L., Laurent, C., Payan, J.-C., & Tisseyre, B. (2022). Observation of shoot growth: A simple and operational decision-making tool for monitoring vine water status in the vineyard. OENO One, 57(1), 235–244. 10.20870/oeno-one.2023.57.1.5481

39. Reynier, A. (2011). *Manuel de viticulture: guide technique du viticulteur*. Lavoisier- Tec&Doc.

40. Songy, A., Fernandez, O., Clément, C., Larignon, P., & Fontaine, F. (2019). Grapevine trunk diseases under thermal and water stresses. Planta, 249(6), 1655–1679. 10.1007/s00425-019-03111-8

41. Spagnolo, A., Larignon, P., Magnin-Robert, M [Maryline], Hovasse, A., Cilindre, C., van Dorsselaer, A., Clément, C., Schaeffer-Reiss, C., & Fontaine, F. (2014). Flowering as the most highly sensitive period of grapevine (Vitis vinifera L. cv Mourvèdre) to the Botryosphaeria dieback agents Neofusicoccum parvum and Diplodia seriata infection. International Journal of Molecular Sciences, 15(6), 9644– 9669. 2014

42. Surico, G., Marchi, G., Mugnai, L., & Braccini, P. (2000). Epidemiology of esca in some vineyards in Tuscany (Italy). Phytopathologia Mediterranea, 39(1), 190–205. 10.1400/57844

43. Surico, G., Mugnai, L., & Marchi, G. (2006). Older and more recent observations on esca: a critical overview.Phytopathologia Mediterranea, 45(4), 68–86. 10.1400/52262

44. Travadon, R., Lecomte, P., Diarra, B., Lawrence, D. P., Renault, D., Ojeda, H., Rey, P [Patrice], & Baumgartner, K. (2016). Grapevine pruning systems and cultivars influence the diversity of wood-colonizing fungi. Fungal Ecology, 24, 82–93. 10.1016/j.funeco.2016.09.003

45. van Niekerk, J., Strever, A. E., Du Toit, G. P., Halleen, F., & Fourie, P. H. (2011). Influence of water stress on Botryosphaeriaceae disease expression in grapevines. Phytopathologia Mediterranea, 50(4), 151–165.

46. Wagschal, I., Abou-Mansour, E., & Petit, A.-N. (2008). 16 Wood diseases of grapevine: A review on eutypa dieback and esca.

