## Supplementary material for "Grapevine vigour: a critical factor driving Trunk Disease expression": Suppl fig 1 and 2

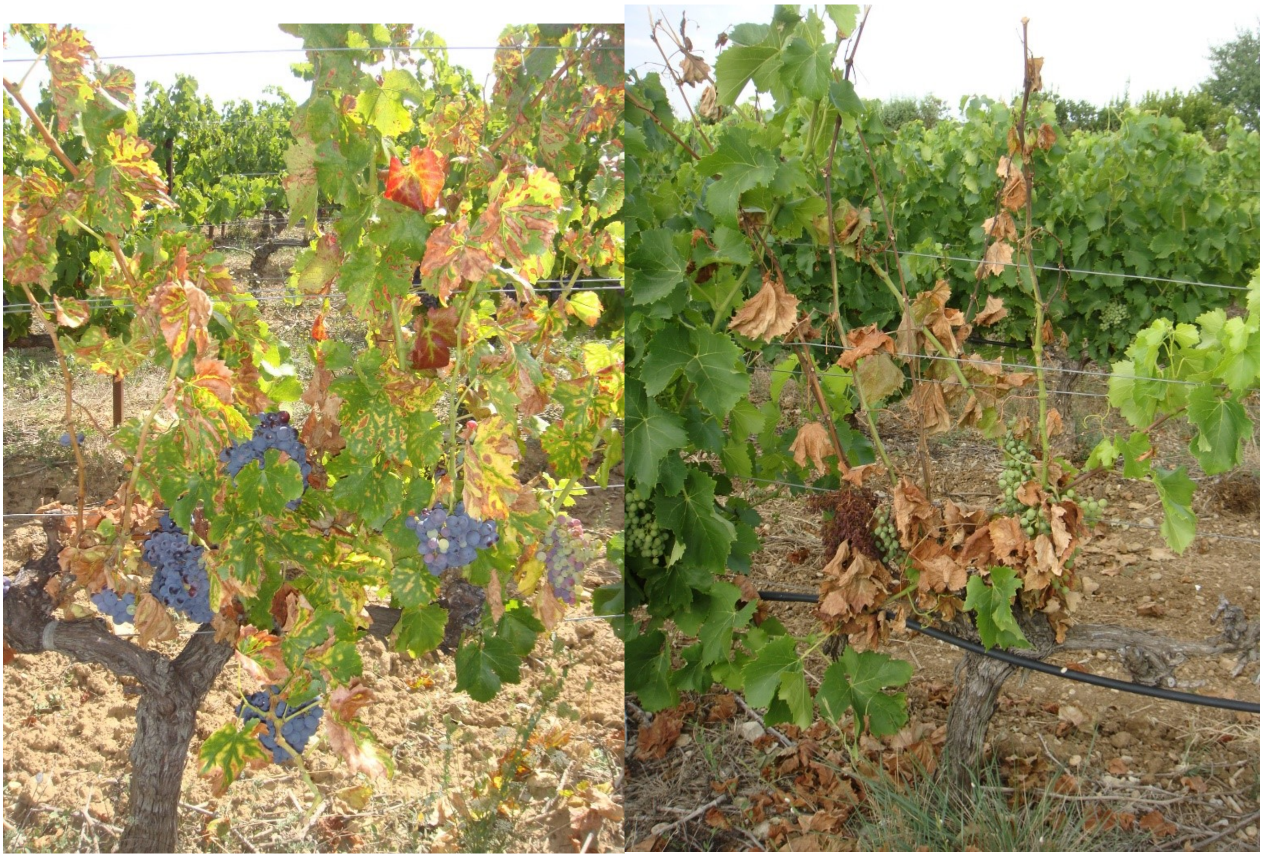

1

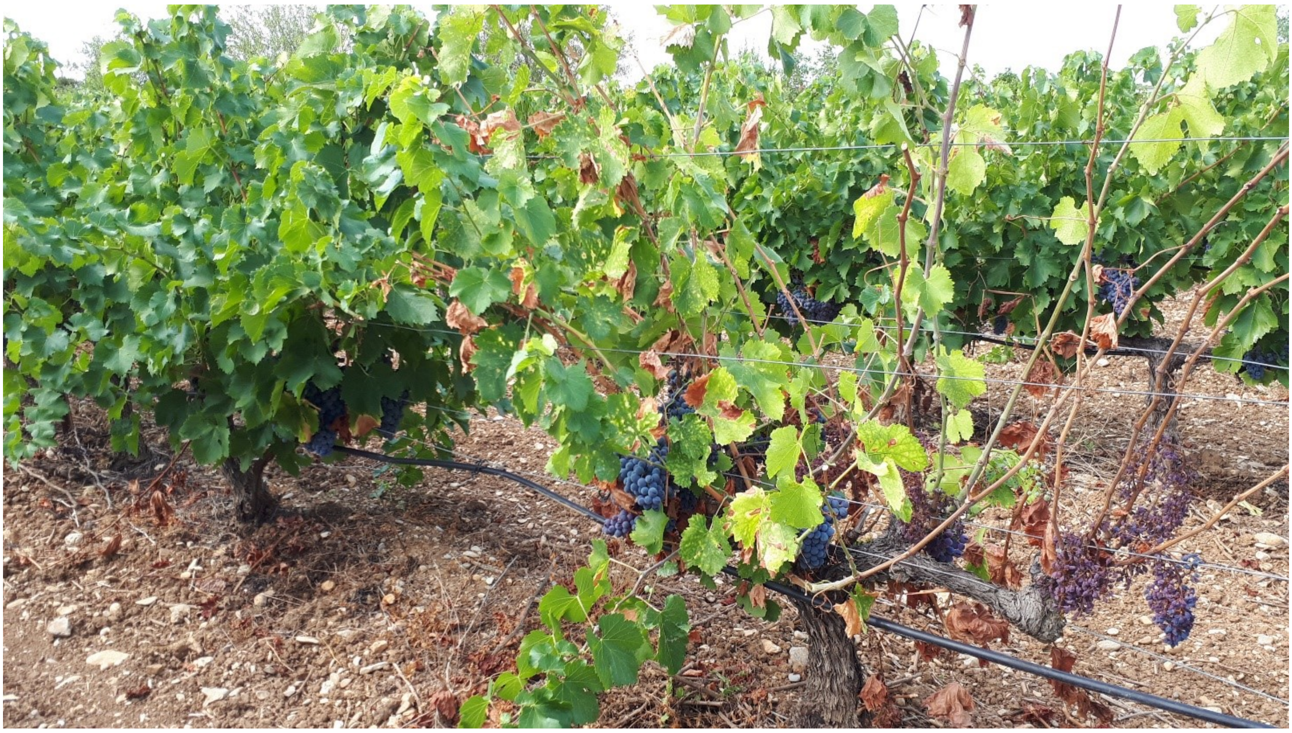

2

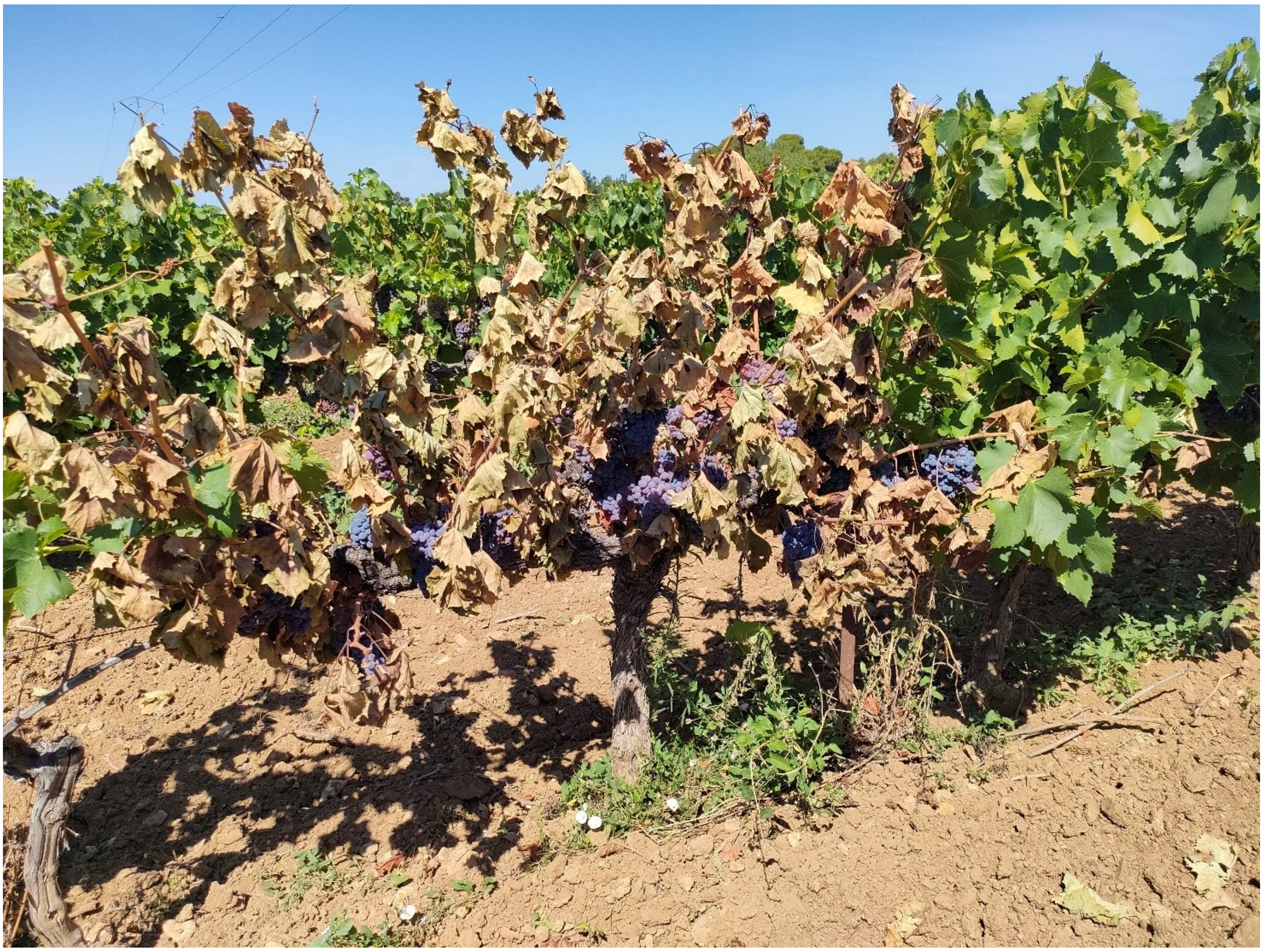

**Supplementary figure 1. Typical expression of EBDA on cultivar Grenache N (a) to (c) typical symptom of severe form (shrivelling or total abscission of the leaves and shoots showing necrotized or desiccated zones alternating with green ones) mixed in greater or lesser proportion with more typical "tiger stripe like" and (d) typical apoplexy form in process. (a) severe form of the entire cep (that will probably be totally desiccated at the end of the season and indistinguishable from apoplexy form), (b) is a milder form with the major proportion of the leaves still present and affected by "tiger stripe-like" symptoms (even if we can still observe missing leaves here and there); (c) is a mix of shoots totally defoliated (and some even dried out) and remaining "tiger stripe-like" affected leaves.**

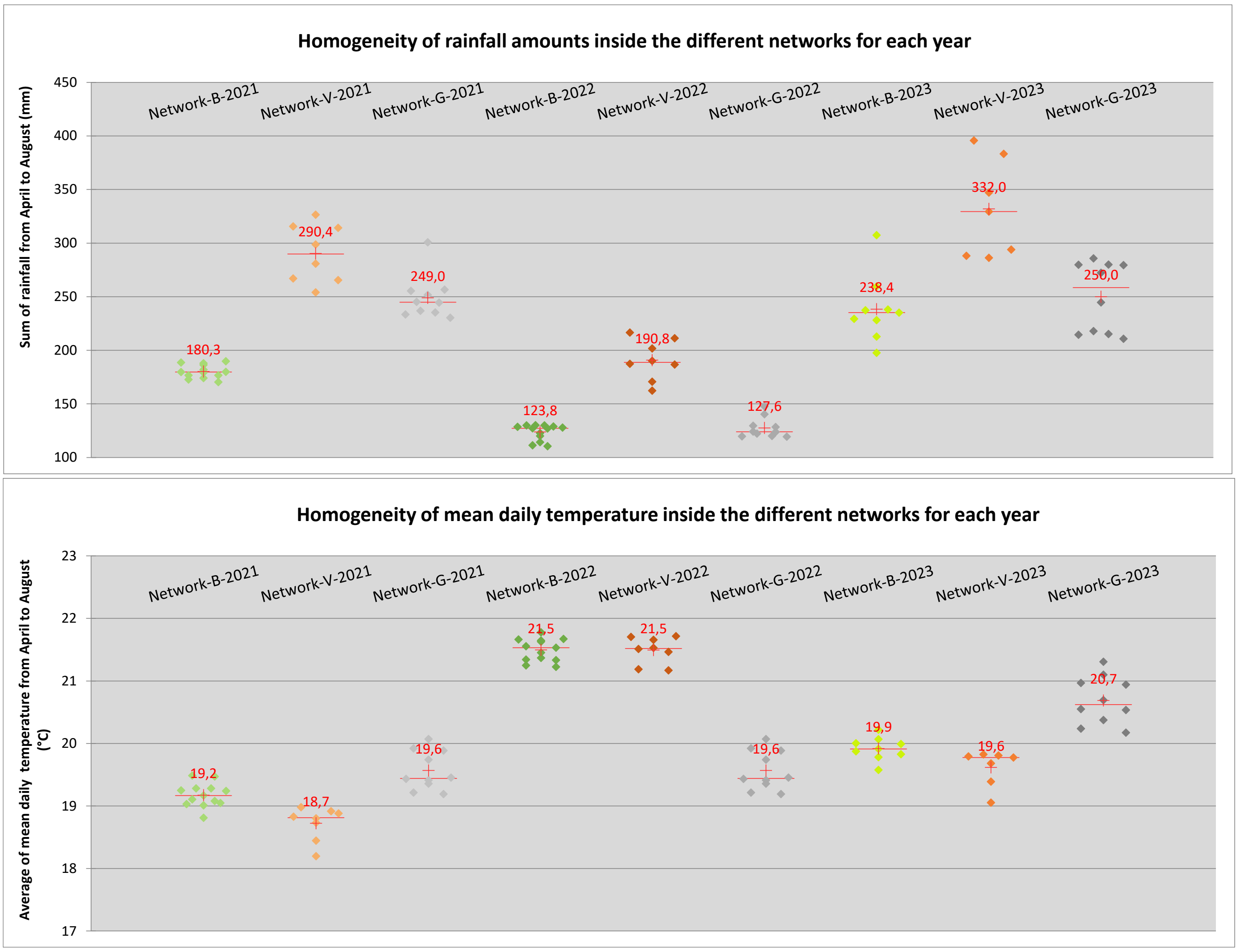


**Supplementary figure 2. Scatter-plot graphs showing the variability of climate for each network and year for (a) rainfall and (b) mean daily temperature.** Each point is a vineyard plot, between 7 and 13 vineyards were chosen per network per year to describe the geographical dispersion of the plots over the area. Red cross is the average value (with red label), red horizontal line is the median value. 2023 is the most heterogeneous year especially for rainfall and in Network-V and Network-G. In Network-V, it corresponds to an east to west gradient of wetness (probably related to the presence of the Ventoux mountain on the eastern side) and in Network-G, least watered area is the NW part of the network.
